# Early-life stromal niches orchestrate B lymphopoiesis at the brain’s borders

**DOI:** 10.1101/2025.11.27.690844

**Authors:** Alec J. Walker, Theodore M. Fisher, Sydney K. Caldwell, Anushree S. Gupte, Justin M. Colville-Reimertz, Bristy Sabikunnahar, Samuel E. Marsh, Yvanka de Soysa, Vahid Gazestani, Alicia C. Walker, Youtong Huang, Ananya V. Mavinkurve, Helena J. Barr, Toby B. Lanser, Sarah Murphy, Sarah Bowling, Lasse Dissing-Olesen, Fernando D. Camargo, Beth Stevens

**Author notes:** These authors contributed equally (Theodore M. Fisher, Sydney K. Caldwell, Anushree S. Gupte).

## Abstract

The dura mater serves as a critical immunological niche for the central nervous system, yet the mechanisms governing the emergence of this niche in early life remain understudied. Here, we chart the trajectory of dural immune development, uncovering a distinctive function for the murine dura as a transient niche for B lymphopoiesis in the early-postnatal window. Shared embryonic progenitors initiate dural B cell development in concert with a multi-organ wave of extramedullary lymphopoiesis that contributes distinctively to the peripheral B cell pool. In the dura, B cells develop locally in discrete sinus-proximal foci, occupying an anatomically defined and developmentally restricted fibroblast niche. Sinus-proximal fibroblasts express the pro-hematopoietic chemokine CXCL12, and local deletion of this crucial factor severely impairs dural B lymphopoiesis. These data reveal a critical function for dural fibroblasts in shaping the early-life B cell compartment and provide a model for how extramedullary niches may support early-life leukocyte production.

## INTRODUCTION

The early-life window (spanning embryonic through early postnatal development) encompasses a period of profound immunological remodeling in mammals, establishing a foundation for life-long immune homeostasis and shaping susceptibility to immune mediated diseases^1–3^. During this period, hematopoietic stem and progenitor cells (HSPCs) migrate to their permanent bone marrow niches, where they fuel production of lymphoid and myeloid cells across the lifespan^4,5^. Shortly thereafter, naïve lymphocytes and antigen presenting cells (APCs) fill secondary lymphoid organs^6–9^, forming the cellular foundation for generation of adaptive immune responses and development of peripheral immune tolerance^10,11^. Importantly, non-lymphoid organs also function as sites for immune cell production and lymphocyte activation during the early-life window^12–18^, highlighting an all-hands-on-deck approach to building the nascent immune system. In primary and secondary lymphoid organs, distinctive stromal cell populations function as professional architects of local immune landscapes^19,20^. However, the cellular and molecular players guiding immune cell development and maturation in other non-lymphoid, early-life organ systems—including within the central nervous system (CNS)—remain poorly defined.

The outermost layer of the meninges—the dura mater—forms a critical immunological niche at the interface between the CNS, the overlying skull, and the peripheral blood, thereby mediating exchange of immune cells and molecules that shape brain health and disease^21,22^. Defining how this immune niche emerges in the early-life window holds broad relevance for understanding brain-immune homeostasis across the lifespan. The adult dura hosts a diverse landscape of lymphoid and myeloid cells predominantly organized around its vascular network^23–25^. These include populations of macrophages that sample brain and blood antigens and protect the CNS from pathogen invasion^25–28^, T cells that modulate brain function and behavior through cytokine signaling^29,30^, and B cells—spanning developmental through plasma cell stages—that support local humoral immunity^31–35^. Dural immune cells intermingle with an intricate network of stromal cells—including fibroblast-like cells (FLCs), pericytes, and smooth muscle cells—that secrete chemokines, growth factors, and extracellular matrix molecules and thereby shape local immune cell function^25,36–38^.

Intriguingly, emerging evidence suggests that some components of the dural immune compartment change markedly across the early-life window, including macrophage and lymphocyte populations that colonize the dura during discrete early-life windows^30,39–41^. However, we lack a comprehensive picture of how the dural immune landscape emerges. Additionally, the stromal players that recruit and maintain immune cells in early-life dural niches—and the functional roles of stromal–immune cell interactions in the early-life dura—remain unexplored.

In this study, we set out to examine the development of the dural immune compartment and to elucidate how early life stromal–immune cell crosstalk governs immune cell colonization and function there. In doing so we uncovered a distinctive function of the early-life dura as a niche for B lymphopoiesis, supporting a transient wave of B cell development spanning the first month of life in mice. To better understand the cellular players that support dural B lymphopoiesis we profiled and functionally interrogated early-life dural stromal cells, uncovering an anatomically and molecularly unique fibroblast niche that critically regulates local B lymphopoiesis via production of the chemokine CXCL12. Together, these findings reveal that brain border tissues contribute to the early-life B cell compartment and establish a framework for studying how non-lymphoid organ stromal niches orchestrate B lymphopoiesis during this critical developmental window.

## RESULTS

### A wave of B cells punctuates the brain border immune landscape in early life

To elucidate how the dural immune compartment emerges in the early-life window, we first aimed to map its composition across postnatal development. We quantified immune cell lineages by flow cytometry at postnatal days (P)1, P6-7, P14, P21, P29-30, and P56-57, spanning developmental milestones including birth (P0), the introduction of solid food (∼P16-17^42^), and weaning (P21 in our facility) (**Figure 1A, Figure S1A-D**). Importantly, we optimized our isolation of the dura to minimize contamination by leukocytes from the adjacent calvarial bone marrow (via a refined dissection protocol, see Methods) and from the blood (via intravenous CD45 labeling and exclusion during flow cytometric analysis) (**Figure S1A-B**). Using this approach, we identified substantial remodeling of the dural immune landscape during the first two months of postnatal life—including distinct patterns of myeloid and lymphoid cell colonization in this window (**Figure 1B-C, Figure S1E-G**). Surprisingly, B cells appeared as a striking developmental wave that peaked at P14 in the dura (**Figure 1B-C**). B cells were largely absent from the dura at P1, increased to nearly 30% of the dural immune compartment by P14, and subsequently diminished by P21 and continued to decrease into adulthood. The magnitude of this B cell wave at P14 was similar in male and female mice (**Figure S1H**) and also appeared in the brain and associated leptomeningeal compartment with similar timing to the dura (**Figure S1I-K**). These data suggest that the first several postnatal weeks represent a critical window for immune cell colonization of the murine dura and pinpoint B cells as a predominant and developmentally regulated feature of this early-life brain border tissue.

**Figure 1.**
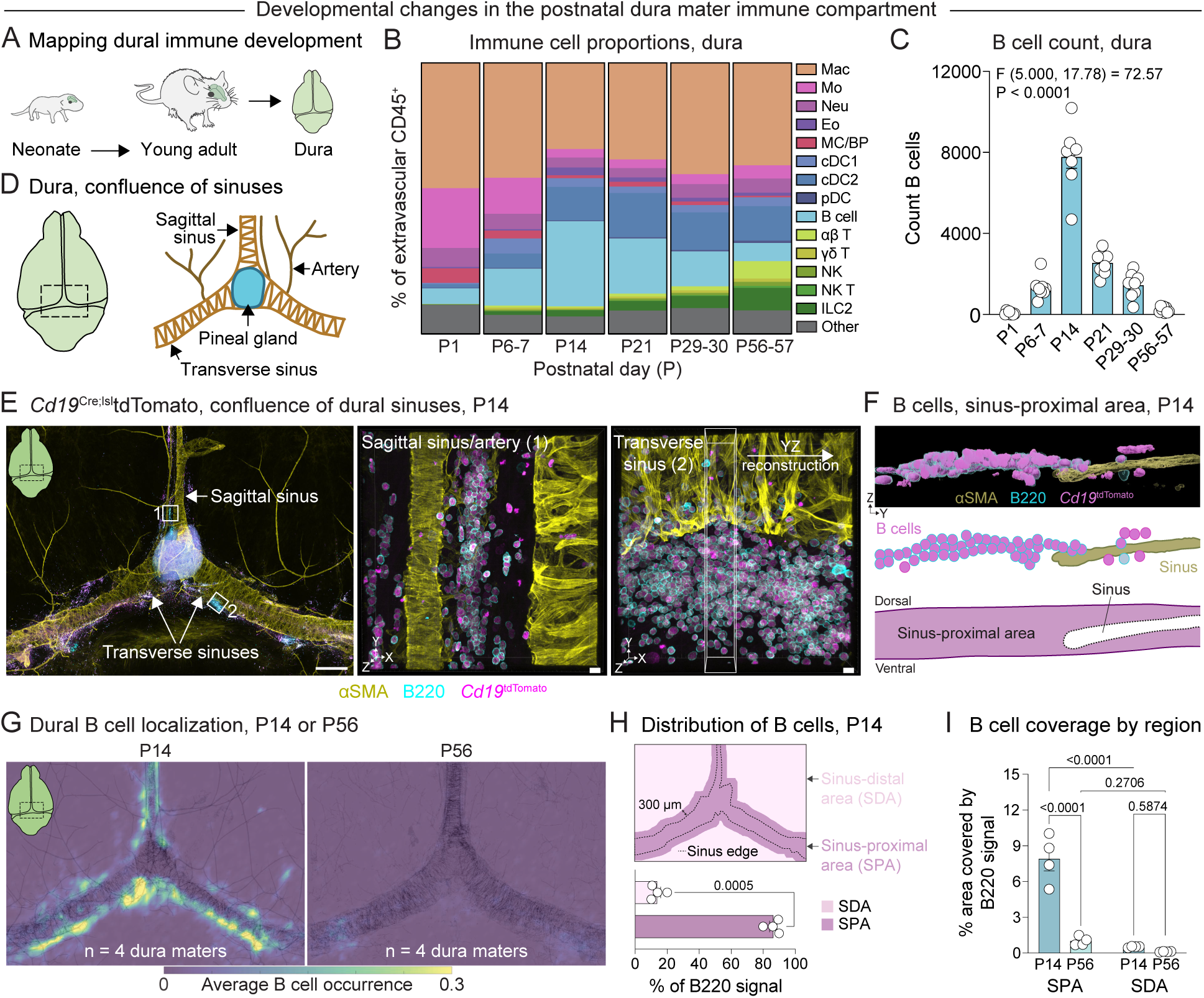
A wave of B cells punctuates the early life dural immune landscape. (A) Experimental schematic-assessing the composition of the dural immune compartment across postnatal development. The dorsal part of the cranial dura was extracted from mice at postnatal days (P)1, P6-7, P14, P21, P29-30, and P56-57 and processed for flow cytometric analysis of immune cell subsets. (B) Quantification of flow cytometry data depicting immune cell proportions in the mouse dura at postnatal developmental timepoints. Each value represents the mean of n = 8 C57BL/6 mice per timepoint collected across two independent experiments for each timepoint. Mac—macrophage; Mo—monocyte; Neu—neutrophil; Eo— eosinophil; MC/BP—mast cell/basophil; cDC1—type 1 conventional dendritic cell; cDC2—type 2 conventional dendritic cell; pDC—plasmacytoid dendritic cell; B cell; αβ T cell; γδ T cell; NK—natural killer cell; NK T—natural killer T cell; ILC2—type 2 innate lymphoid cell; Other—uncategorized CD45^+^ cells. (C) Quantification of flow cytometry data depicting total CD19^+^ B cell counts in the mouse dura at postnatal developmental timepoints. n = 8 mice C57BL/6 mice per timepoint collected across two independent experiments for each timepoint. Error bars represent SEM. Welch’s ANOVA. (D) Schematic depicting a whole mount preparation of the mouse dura mater and attached pineal gland. The area near the confluence of the sagittal and transverse sinuses is highlighted. (E) Representative tile scan image of the confluence of the dural sinuses in a P14 *Cd19*^Cre^;^lsl^tdTomato mouse (left). Imaris representations of high magnification images depict B cell clusters near the sagittal sinus (region of interest 1, middle) and transverse sinus (region of interest 2, right). The approximate locations of the high magnification images are indicated on the tile scan with white boxes. `α`SMA—`α`-smooth muscle actin, yellow; B220—B cells, cyan; *Cd19*^tdTomato^—B cells, magenta. A segment of the image depicting the transverse sinus was reconstructed in 3-dimensions and is shown in (F). Scale bar for tile scan image indicates 500 µm. Scale bars for high magnification images indicate 10 µm. (F) Imaris surface rendering of a B cell cluster near the transverse dural sinus at P14 (from panel E) shown in the YZ plane (top), and corresponding diagram depicting the spatial relationship between B cells and the transverse sinus (middle). B cells at P14 are frequently located in the region adjacent to the dural sinuses, here referred to as the sinus-proximal area (purple highlighted area located dorsal to, ventral to, and with 300 µm of the lateral edge of the sinuses, bottom diagram). (G) Spatial localization of B cells in the dura of P14 and P56 mice. n = 4 mice per age merged to create one heatmap that represents the average occurrence of B220 signal at each given XY coordinate across the dura. Heatmaps are overlaid with a composite image of aSMA signal (black outline) from the same four mice at each age. (H) Spatial distribution of B cells (B220 signal) in different regions of the dura, including the region directly overlying and within 300 µm of the outer edge of the sinuses (sinus-proximal area, SPA), and the region more than 300 µm from the edges of the sinuses (sinus-distal area, SDA). n = 4 mice, using data from the same mice employed to generate the P14 heatmap in panel (G) except with quantifications performed on a per mouse basis. Error bars represent SEM. Student’s T-test, paired, two-tailed. (I) Percent area of each dural region that is covered by B cells (B220 signal). n = 4 mice per age, using data from the same mice employed to generate the heatmaps in panel (G), except with quantifications performed on a per mouse basis. SPA—sinus-proximal area; SDA—sinus-distal area. Error bars represent SEM. Two-way repeated measures ANOVA with Uncorrected Fisher’s LSD. Interaction: F (1, 6) = 39.35, P = 0.0008.

To examine the spatial localization of early-life B cells, we prepared whole mount dural samples from *Cd19*^Cre^;^lsl^tdTomato mice at P14. We found that tdTomato^+^B220^+^ B cells formed large clusters predominantly near the transverse sinuses, the caudal portion of the sagittal sinus, the confluence of the sinuses, and arteries and veins near the confluence of the sinuses (**Figure 1D-E, Figure S2A-B**), prompting us to focus on these regions for further analyses. B cell clusters also appeared at the rostral-rhinal hub^35^ but were largely absent from other rostral areas of the dura or perivascular regions distant from the sinuses (**Figure S2B**). Further examination of B cell clusters in three dimensions revealed their localization to a unique microanatomical region adjacent to the dural sinuses, which we define here as the sinus-proximal area (**Figure 1F**). We also examined the immune composition of early-life dural B cell clusters, hypothesizing that they might represent tertiary lymphoid follicles like those found in the early-life lung and intestines^16,43,44^ or in the aged/inflamed dura^35^. However, sinus-proximal B cell clusters contained few CD3^+^ T cells (**Figure S2C**), arguing against this possibility and instead suggesting a different function for these early-life immune structures compared to those observed in the adult dura.

To confirm the developmental kinetics of the B cell wave using an orthogonal approach, we constructed heatmaps of B cell localization across multiple dural samples at P14 and P56 using a custom MATLAB pipeline (**Figure S2D-E**). Additionally, we quantified B cell occupancy of the sinus-proximal area (overlying and within 300 μm of the sinuses) and the sinus-distal area (>300 μm from the sinuses) (**Figure S2F)**. These analyses demonstrated that B cells consistently and preferentially localized to the sinus-proximal area at P14 and revealed a marked decrease in B cell coverage of this area from P14 to P56 (**Figure 1G-I**). Together, these data suggest that the sinus-proximal area of the dura may represent a unique microenvironment that is exquisitely tuned to support B cells in the early-life window.

### Early-life B cells undergo lymphopoiesis in brain border tissues

To elucidate the phenotype of B cells present in the dura and other early-life CNS tissues, we performed single-cell RNA sequencing (scRNA-seq) on CD45^+^ immune cells from the dura and the pooled brain and leptomeninges (brain-leptomeninges) of P14 mice (**Figure S3A**). We also included CD45^+^ cells from the calvarial bone marrow and tibial bone marrow, which allowed us to compare CNS B cells to abundant populations of early-life bone marrow B cells. Finally, we included CD45^-^TER119^-^ non-immune cells from the dura, calvarial bone marrow, and tibial bone marrow to capture stromal and endothelial niche cells that likely govern B cell recruitment to and maintenance in these early-life organs^19^.

Initial clustering of this data revealed diverse populations of immune cells, stromal cells, endothelial cells, and other non-immune cell types across the tissues sampled (**Figure S3B-G**). B cells represented a substantial fraction of the dural and brain-leptomeningeal immune compartments and closely overlapped in UMAP space with B cells from the calvarial and tibial bone marrow (**Figure 2A-B, Figure S3F).** Sub-clustering of the B cell compartment revealed that dural and brain-leptomeningeal B cells (hereinafter collectively CNS B cells) recapitulated stages of B lymphopoiesis, including Pro, Pre, and newly formed (immature/transitional) B cells (**Figure 2C-E**). Pro and Pre B cells from CNS tissues expressed the early B cell marker *Cd93*, the B cell growth factor receptor *Il7r*, and the chemokine receptor *Cxcr4*, whereas newly formed B cells showed elevated expression of *Ms4a1* (CD20) and the B cell receptor (BCR) heavy chain gene, *Ighm* (IgM) (**Figure 2F-G**). Mature B cells—enriched for expression of *Ighd* (IgD), *H2-Ab1*, and *Ms4a4c*^45^—also appeared in all compartments, though they represented a minority of B cells as did plasma cells expressing *Jchain* and *Ighm*.

**Figure 2.**
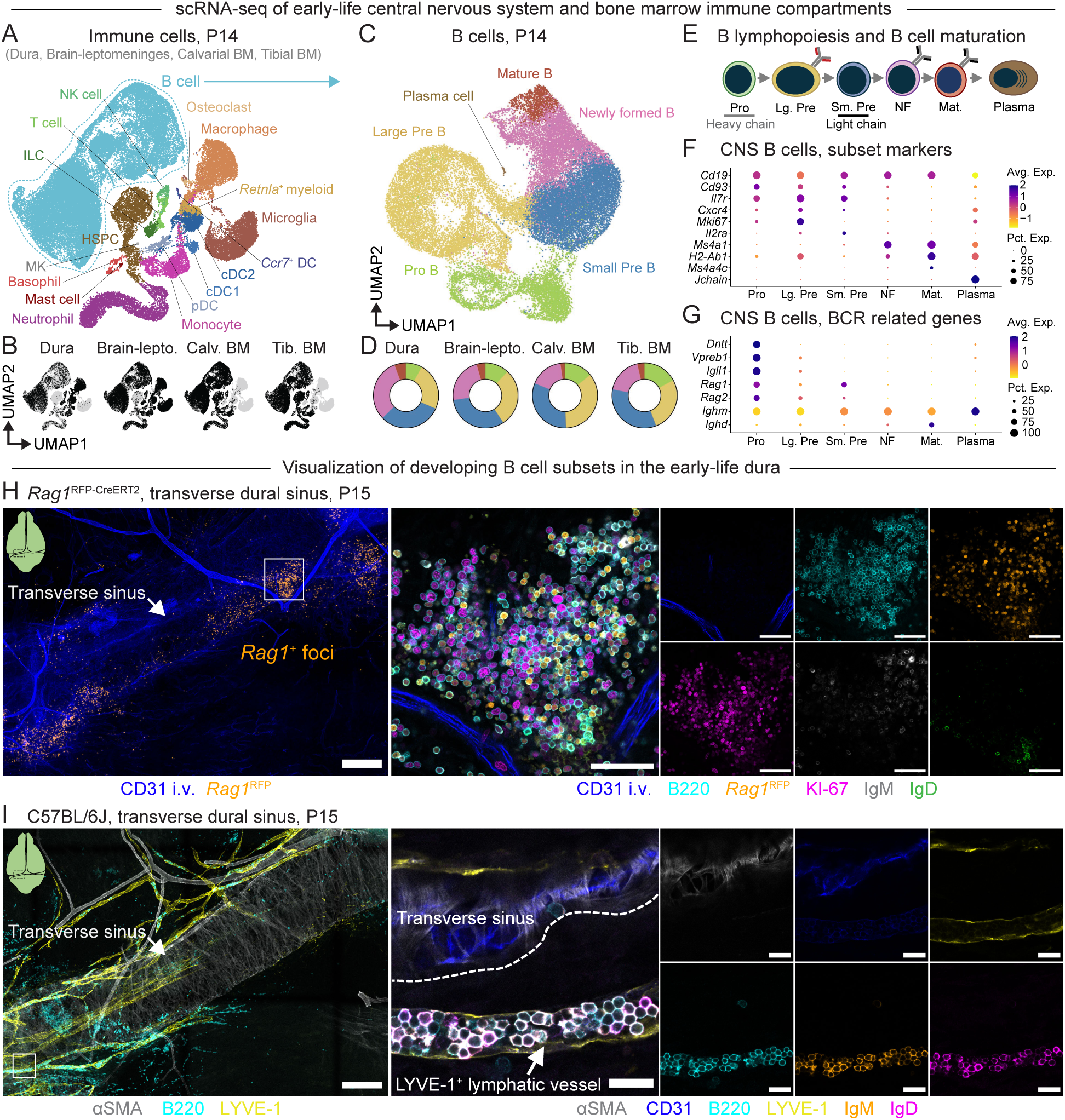
Early-life B cells undergo lymphopoiesis in brain border tissues. (A) UMAP representation of scRNA-seq data depicting immune cells from dura, pooled brain and leptomeninges, calvarial bone marrow, and tibial bone marrow at P14. B cells are highlighted by a dotted lasso. n = 3 male C57BL/6 mice. cDC1—type 1 conventional dendritic cell; cDC2—type 2 conventional dendritic cell; pDC—plasmacytoid dendritic cell; *Ccr7*^+^ DC—*Ccr7*^+^ dendritic cell; NK cell—natural killer cell; ILC—innate lymphoid cell; HSPC— hematopoietic stem and progenitor cell; MK—megakaryocyte. (B) Distribution of immune cells in each organ. Brain-lepto.—pooled brain and leptomeninges; Calv. BM—calvarial bone marrow; Tib. BM—tibial bone marrow. (C) UMAP representation of B cells across dura, pooled brain and leptomeninges, calvarial bone marrow, and tibial bone marrow compartments. B cells are grouped by subtype, predominantly representing different stages of B lymphopoiesis. (D) Proportion of B cell subtypes in each organ, related to (C). Brain-lepto.—pooled brain and leptomeninges; Calv. BM—calvarial bone marrow; Tib. BM—tibial bone marrow. (E) Schematic—stages of B lymphopoiesis and B cell maturation. Recombination of the BCR heavy chain locus occurs at the Pro B cell stage and recombination of light chain locus occurs at the small Pre B cell stage. Pro B cell, large (Lg.) Pre B cell, small (Sm.) Pre B cell, newly formed (NF) B cell, mature (Mat.) B cell, plasma cell. Newly formed B cells include immature and transitional B cell subtypes in this representation. (F) Expression of canonical genes associated with subtypes of early and mature B cells, plotted for dura and brain-leptomeninges B cells (collectively, CNS B cells). Pro B cell, large (Lg.) Pre B cell, small (Sm.) Pre B cell, newly formed (NF) B cell, mature (Mat.) B cell, plasma cell. (G) Expression of genes related to BCR recombination, plotted for dura and pooled brain and leptomeninges B cells (collectively, CNS B cells). Pro B cell, large (Lg.) Pre B cell, small (Sm.) Pre B cell, newly formed (NF) B cell, mature (Mat.) B cell, plasma cell. (H) Representative tile scan image (left) and high magnification images (right) of the transverse sinus of a *Rag1*^RFP-CreERT2^ dura at P15 depicting foci of *Rag1*^+^ B cells. Intravenous (i.v.) CD31—vasculature, blue; B220—B cells, cyan; RFP—*Rag1*^+^ cells; orange; KI-67—proliferating cells, magenta; IgM, white; IgD, green. The approximate location of the high magnification image is indicated with a white box on the tile scan image. Scale bar for tile scan image indicates 200 µm. Scale bars for high magnification images indicate 50 µm. (I) Representative tile scan image (left) and high magnification images (right) of the transverse sinus of a C57BL/6 dura at P15 depicting B cell association with sinus-proximal lymphatic vasculature. `α`SMA—α-smooth muscle actin, white; CD31—vasculature, blue; LYVE1—lymphatic vasculature, yellow; B220—B cells, cyan; IgM, orange; IgD, magenta. The approximate location of the high magnification image is indicated with a white box on the tile scan image. Scale bar for tile scan image indicates 200 µm. Scale bars for high magnification images indicate 20 µm.

As B cells differentiate from hematopoietic precursors into their mature form, they undergo recombination of the heavy and light chain regions of the BCR locus in Pro and small Pre B cells, respectively (**Figure 2E**). Indeed, in our dataset, Pro B and Pre B cells from CNS tissues expressed the DNA recombinases *Rag1* and *Rag2* (**Figure 2G**), which together with DNA repair machinery mediate BCR recombination^46^. Pro B cells also expressed terminal deoxynucleotide transferase (TdT, *Dntt*) and components of the surrogate light chain, *Vpreb1* and *Igll1* (**Figure 2G**), which mediate diversification and stabilization of the BCR heavy chain, respectively^47,48^. To investigate the spatial organization of these developing B cell subtypes in the early-life dura, we took advantage of mice that express RFP downstream of the *Rag1* promoter (*Rag1*^RFP-CreERT2^). At P15, we observed distinct lymphopoietic foci containing *Rag1*^+^IgM^-^IgD^-^ Pro/small Pre B cells located near the dural sinuses (**Figure 2H**). Dural lymphopoietic foci also contained Rag1^lo/-^KI-67^+^IgM^-^IgD^-^ large Pre B cells and to a lesser extent newly formed B cells expressing membrane-associated IgM and/or IgD. Interestingly, a subset of IgM^+^ and/or IgD^+^ B cells (likely newly formed B cells) appeared tightly packed inside of lymphatic vessels that ran alongside the transverse sinuses (**Figure 2I**). Together, these data indicate that early life dural B cell development occurs in distinct lymphopoietic foci proximal to the dural sinuses and nominate lymphatic vasculature as one potential route of egress for CNS-born B cells as this early life lymphopoietic wave subsides.

### Dural B cells compose part of an early life lymphopoietic wave that contributes distinctively to the mature B cell pool

While the bone marrow functions as the predominant site of murine B lymphopoiesis in adulthood^5^, other organs including the mouse spleen and small intestine lamina propria contain developing B cells in the perinatal window^12,49,50^. To assess whether the wave of dural B lymphopoiesis coincides with B cell development in other early-life organs, we quantified developing B cells in primary lymphoid organs (liver, tibial bone marrow, calvarial bone marrow) and extramedullary organs including CNS tissues (dura, brain-leptomeninges, spleen, small intestine and Peyer’s patches), at P7, P14, P21-22, P27-28, and P34-36 by flow cytometry (**Figure 3A, Figure S4A-B**).

**Figure 3.**
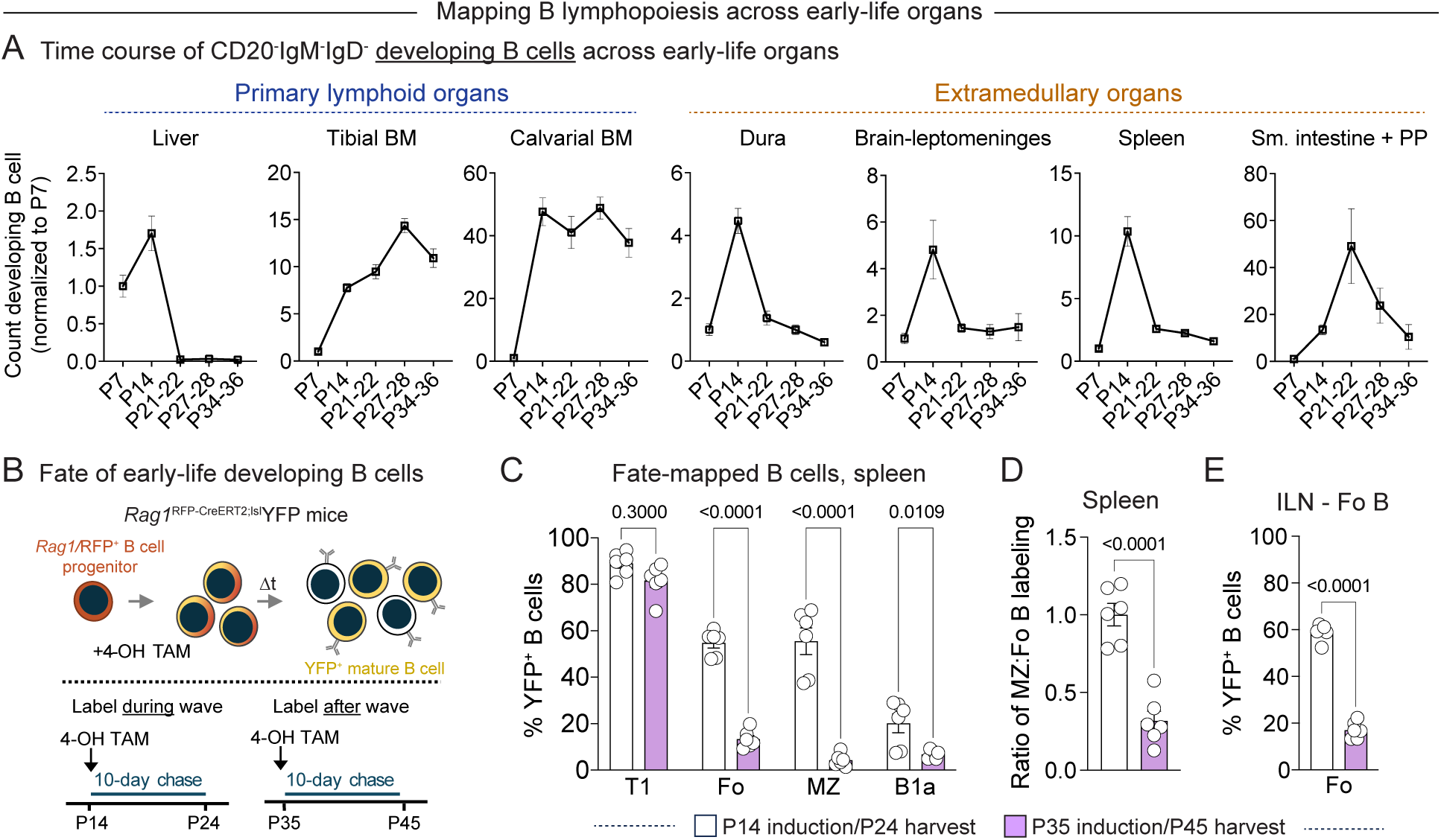
A multi-organ, early life lymphopoietic wave contributes distinctively to the peripheral B cell pool. (A) Normalized count of developing B cells from primary lymphoid and extramedullary organs across a time course of postnatal development, assessed by flow cytometry. Numbers of B cells for each organ were normalized to the mean B cell count of that organ at P7. n = 8 C57BL/6 mice per age at P7, P27-28, and P34-36, and 12 mice per age at P14 and P21-22, pooled from 2-3 independent experiments per age. Error bars represent SEM. Welch’s ANOVA. Liver: F(4.000, 19.37) = 23.82, P < 0.0001; Tibial bone marrow (BM): F(4.000, 18.79) = 132.1, P < 0.0001; Calvarial bone marrow: F(4.000, 17.25) = 93.98, P < 0.0001; Dura: F(4.000, 20.17) = 22.83, P < 0.0001; Brain-leptomeninges: F(4.000, 19.39) = 2.356, P = 0.0896; Spleen: F(4.000, 20.76) = 24.53, P < 0.0001; Small intestine + Peyer’s patches (PP): F(4.000, 17.28) = 12.93, P < 0.0001. (B) Fate mapping of early-life *Rag1*^+^ B cells. *Rag1*^RFP-CreERT2;lsl^YFP mice were pulse labeled at induction timepoints P14 or P35, corresponding to the peak of the early life lymphopoietic wave or after the wave subsides. Transitional and mature B cell populations in the spleen and inguinal lymph nodes were evaluated for YFP labeling ten days later at the indicated harvest timepoint. (C) Percent of splenic transitional T1, follicular (Fo), marginal zone (MZ), and B1a B cell populations expressing YFP, indicating fate mapping from *Rag1*^+^ cells. n = 6 mice for each induction timepoint pooled from two independent experiments per induction timepoint. Error bars represent SEM. Two-way ANOVA with Sidak’s multiple comparisons test. Timepoint-B cell subset interaction: F(3, 40) = 25.53, P < 0.0001. Select post-hoc comparisons are shown. (D) Ratio of the percent of splenic marginal zone B cells labeled with YFP to the percent of follicular B cells labeled with YFP in *Rag1*^RFP-CreERT2;lsl^YFP mice induced at P14 or P35. n = 6 mice for each induction timepoint pooled from two independent experiments per induction timepoint. Error bars represent SEM. Student’s unpaired T test, two-tailed. (E) Percentage of inguinal lymph node (ILN) follicular (Fo) B cells expressing YFP, indicating fate mapping from *Rag1*^+^ cells. n = 6 mice for each induction timepoint pooled from two independent experiments per induction timepoint. Error bars represent SEM. Student’s unpaired T test, two-tailed.

As expected, we found that B lymphopoiesis in the liver declined, and B lymphopoiesis in the tibial bone marrow rapidly increased, from P7 to P28 (**Figure 3A**), reflecting the perinatal transition from fetal liver to bone marrow blood cell production in mice^51^. In contrast, numbers of developing B cells in the dura, brain-leptomeninges, and spleen peaked at P14 and rapidly declined through P21 and P28 timepoints. Numbers of developing B cells also peaked in the small intestine and Peyer’s patches during this early-life window, though this peak was shifted toward P21 as previously described^12^. In contrast, numbers of CD20^+^ (newly formed and mature) B cells gradually increased with age in most organs, except for in the dural and the brain-leptomeningeal compartments where their trajectories closely mirrored those of developing B cells (**Figure S4C**). Thus, dural B lymphopoiesis composes part of a coordinated wave of B cell development occurring across multiple extramedullary tissues in the early-life mouse.

Given this data, we investigated how B cells born during this early-life wave contribute to the postnatal B cell pool. To compare the fate of B cells derived from this wave to those derived from a young adult timepoint, we pulse labeled *Rag1*^RFP-CreERT2^;^lsl^YFP mice with 4-hydroxytamoxifen (4-OH TAM) at P14 (the peak of the lymphopoietic wave) or P35 (after the wave) (**Figure 3B**). We reasoned that this approach would label *Rag1*^+^ developing B cells present during each of these distinct time windows and allow us to identify their YFP^+^ progeny as they differentiate into mature B cell subsets including B1 B cells, follicular B cells, and marginal zone B cells^52^. Ten days after labeling, we harvested organs that host mature B cell populations—including the spleen and inguinal lymph nodes (ILN)—and assessed the contribution of fate-mapped B cells to these compartments (**Figure 3B, Figure S4D**). Induction at P14 or P35 led to robust and equivalent labeling of the T1 transitional B cell compartment in the spleen, indicating that these recent splenic immigrants had differentiated from *Rag1*-expressing cells within this experimental time window (**Figure 3C**). *Rag1*^+^ B cells pulse-labeled at P14 additionally contributed to the splenic B2 B cell compartment by harvest timepoint P24, generating approximately 50-60% of follicular B cells and 40-70% of marginal zone B cells present at this time. Induction at P14 also led to some labeling of the splenic B1a B cell compartment, though this labeling was lower than that observed in the B2 B cell compartment. In contrast, *Rag1*^+^ B cells pulse labeled at P35 contributed substantially less to the follicular B cell compartment (∼10-20%) and less yet to the marginal zone B cell compartment (∼1-9%) ten days after induction (**Figure 3C**). Indeed, the ratio of YFP-expressing cells in the marginal zone B cell compartment compared to the follicular B cell compartment decreased markedly between P14 and P35 induction timepoints (**Figure 3D**), suggesting that the relative output of marginal zone B cells—or their specification in the periphery^53^—diminishes after this early-life lymphopoietic wave subsides. We observed similarly high contribution of *Rag1*^+^ B cells labeled at P14 to follicular B cells in the ILNs and substantial loss of this contribution after induction at P35 (**Figure 3E**). This data suggests that the early-life wave of B lymphopoiesis contributes to rapid generation of the B2 B cell compartment and furnishes the spleen with marginal zone B cells whose replacement by B cells from other sources may be limited in older mice.

### A shared pool of progenitors initiates medullary and extramedullary B lymphopoiesis

Given the similarly timed initiation of B lymphopoiesis in early-life organs, we next sought to assess how this process is coordinated across the body. We hypothesized that there must exist a common migration event that initially seeds different early-life organs with B cell progenitors or early B cells, perhaps corresponding to the well-described transition from fetal liver to bone marrow hematopoiesis^51^. To investigate the origins of developing B cells in the dura and other early-life B cell niches, we leveraged the CRISPR array repair lineage tracing (CARLIN) system^54^. CARLIN;M2-rtTA;TetO-Cas9 mice allow Cas9-mediated recombination of a DNA barcode locus after induction with doxycycline *in vivo*, generating unique barcodes that are heritable, transcribed into mRNA, and can be read out by RNA sequencing. We reasoned that—by inducing CARLIN barcode recombination at different embryonic and early postnatal timepoints—we could label early-life HSPCs and subsequently assess the lineage relationships between their B cell progeny that contribute to this postnatal lymphopoietic wave (**Figure 4A**). We chose three induction timepoints: embryonic day (E)12.5 (when HSPCs reside predominantly in the fetal liver); E17.5 (shortly after the onset of HSPC migration to the bone marrow and spleen); and P3 (when substantial pools of HSPCs have colonized the bone marrow and spleen)^4,51^. For each induction timepoint, we sorted developing B cells (pooled Pro and Pre B cells) from the dura, calvarial bone marrow, leg (tibial and femoral) bone marrow, and spleen at P13-P15 for targeted RNA sequencing of the CARLIN barcode locus (**Figure 4A, Figure S5A**).

**Figure 4.**
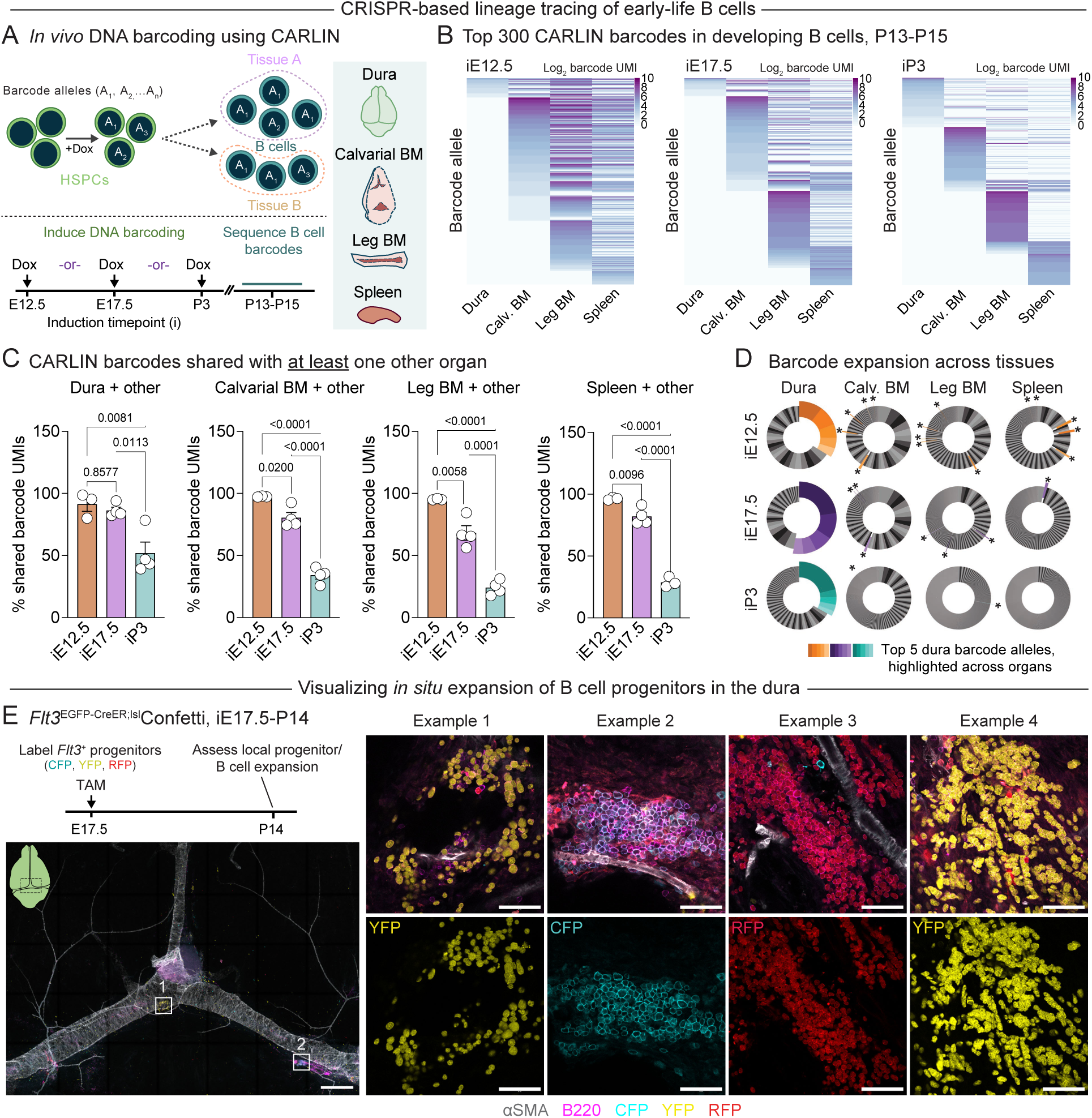
A shared pool of progenitors initiates early life B lymphopoiesis and seeds defined lymphopoietic foci in the dura. (A) Experimental design for DNA barcode-based lineage tracing of early-life B cells using CARLIN. CARLIN mice express a DNA barcode locus that is recombined by doxycycline-inducible Cas9 to generate unique, heritable barcodes that can be read out by bulk RNA sequencing. Doxycycline (Dox) was administered to CARLIN;M2-rtTA;Tet-O-Cas9 mice at different embryonic and postnatal induction timepoints, allowing recombination of barcodes in HSPCs that could then be traced into resultant B cell populations in the dura, calvarial bone marrow (BM), leg (tibial + femoral) bone marrow, and spleen at P13-P15. (B) Representative heatmaps displaying the top 300 CARLIN barcode alleles found in developing B cells (pooled Pro and Pre B cells) from dura, calvarial bone marrow, leg (tibial + femoral) bone marrow, and spleen in P13-P15 mice where CARLIN barcoding was induced at the indicated developmental timepoints. The count of unique motif identifiers (UMIs) for each barcode allele is plotted on a log_2_ scale, correlating with the number of B cells expressing each barcode allele. (C) Quantification of CARLIN barcode sharing between early life organs, displaying the percentage of CARLIN barcode UMIs that share an allele sequence with barcodes in at least one other organ. n = 3 mice at iE12.5, 4 mice iE17.5, and 4 mice at iP3. Error bars represent SEM. One-way ANOVA with Tukey’s multiple comparisons test. Dura: F(2, 8) = 11.07, P = 0.0050; calvarial bone marrow: F(2, 8) = 95.91, P < 0.0001; leg bone marrow: F(2, 8) = 70.15, P < 0.0001; spleen: F(2, 8) = 212.3, P < 0.0001. (D) Clonal expansion of CARLIN barcode alleles in each organ from one representative mouse per induction timepoint. The top 5 expanded dural clones at each induction timepoint are highlighted to facilitate comparison across organs in the same mouse. Asterixis around the perimeter of the calvarial bone marrow (Calv. BM), leg bone marrow, and spleen plots indicate barcode alleles corresponding to the top 5 expanded barcode alleles in the dura. (E) Experimental design for evaluating clonal expansion of B cell progenitors *in situ*. *Flt3*^EGFP-CreER^;^lsl^Confetti mice were induced with tamoxifen (TAM) at iE17.5 and the dura imaged at P14 to assess the spatial relationship between B cells derived from distinct (cyan, yellow, or red fluorescent protein-expressing) progenitor clones. Representative tile scan image (left) and high magnification images (right) of whole mount dura from P14 *Flt3*^EGFP-CreER^;^lsl^Confetti mice. aSMA—a-smooth muscle actin, white; B220—B cells, magenta; CFP, YFP, and RFP—Confetti clones. GFP clones were not examined as both the *Flt3*^EGFP-CreER^ and the Confetti mice contain a GFP allele. Tile scan is displayed as a maximum intensity projection of multiple Z-stacks, scale bar 500 µm. High magnification images represent single plane confocal images of B cell clusters, scale bar 50 µm. Boxed insets on the tile scan image indicate the approximate location of high magnification images “Example 1” and “Example 2.”

This data revealed distinct patterns of B cell barcode distribution across organs that were dependent on the induction timepoint (**Figure 4B**). After barcode labeling at induction timepoint (i)E12.5, almost all CARLIN barcode counts (unique motif identifiers - UMIs) found in dural B cells shared an allele sequence with barcodes found in B cells from another organ (**Figure 4C**). This pattern also held true when comparing barcodes in B cells from the calvarial bone marrow, leg bone marrow, or spleen with those found in other organs, suggesting a common embryonic origin of dural and other early-life developing B cells. B cells from mice induced at iE17.5 showed slightly diminished barcode sharing across organs compared to iE12.5. However, induction at iP3 lead to a substantial loss of multi-organ barcode sharing compared to earlier timepoints (**Figure 4C**). Together, these results support a model where a common pool of embryonic HSPCs or early B cell progenitors migrate to medullary and extramedullary organs—including the dura—in the perinatal window to seed coordinated waves of B lymphopoiesis.

We also examined pairwise B cell barcode sharing between the dura and other individual organs. The total fraction of dural B cell barcode UMIs that shared an allele sequence with B cell barcodes in the calvarial bone marrow, leg bone marrow, or spleen each decreased from iE12.5 to iP3 labeling timepoints (**Figure S5B**). Dural and calvarial bone marrow B cells also showed an increase in the percentage of barcode UMIs that were exclusively shared between themselves—but not with other organs—after labeling at iP3 compared to labeling at iE12.5 (**Figure S5C**). In contrast, dural B cells showed no change in exclusive barcode sharing with the leg bone marrow or spleen across induction timepoints. These data suggest that a small pool of B cells or their progenitors continue to migrate between the dura and the calvarial bone marrow even after systemic migration of progenitors diminishes in the perinatal period, consistent with previous reports of leukocyte exchange between these compartments^22,31,55–57^.

### B cell progenitors expand locally in microanatomical dural niches

To explore the dynamics of early B cell expansion in different organs, we also examined the clonality of B cell barcode alleles. In the dura, the top five expanded CARLIN barcode alleles comprised a substantial fraction of total barcode UMIs within the B cell pool (**Figure 4D**). However, the top expanded dural barcode alleles generally did not overlap with the top expanded alleles in other organs, indicating that these clones expanded locally after migration to the dura. Since we specifically analyzed developing B cells that lacked surface expression of the BCR for these experiments, it is likely that BCR-independent proliferation of B cell progenitors mediated this expansion. Thus, these data suggest that early-life dural B cells arise from common embryonic progenitors as do early-life bone marrow and splenic B cells but subsequently diverge and expand locally to generate a largely independent wave of B cell development.

Given the substantial clonal expansion of CARLIN barcodes we observed at P13-P15, we also wanted to visualize the spatial relationship between dural B cells derived from distinct progenitor clones. We first used *Flt3*^EGFP-CreER^;^lsl^tdTomato mice to determine the developmental timing with which *Flt3*-expressing progenitors (i.e. multipotent progenitors (MPP), common lymphoid progenitors (CLP), or Pre-Pro B cells) differentiate into B cells in the perinatal window (**Figure S5D-E)**. Pulse labeling with tamoxifen at iE17.5 led to near-complete labeling of MPPs in the bone marrow and spleen, and of developing B cells in all organs examined, at P14 (**Figure S5F-H**). Tamoxifen induction at iP3 or iP7 lead to consistently high labeling of MPPs but decreased labeling of B cells at P14, suggesting that FLT3^+^ progenitors differentiate into B cells during the perinatal window to fuel early-life lymphopoiesis in the dura and other organs. We next used *Flt3*^EGFP-CreER^;^lsl^Confetti mice to examine the spatial dynamics of B cell progenitor expansion in the dura during this same window. We pulse labeled these mice at iE17.5 with tamoxifen, driving heritable expression of one of four fluorescent proteins—CFP, YFP, RFP, and GFP (not assessed due to overlap with *Flt3*-driven EGFP)—in *Flt3*^+^ progenitors (**Figure 4E**). Imaging of whole mount dural preparations from these mice at P14 revealed topographically defined clusters of B cells, most of which were each predominated by a single Confetti clone (**Figure 4E**). Together with our CARLIN data demonstrating substantial expansion of select dural B cell barcode alleles, these results suggest that B cell progenitors colonize microanatomical niches in the perinatal dura and expand extensively *in situ* to generate this early life lymphopoietic wave.

### Early-life dural and bone marrow stromal niches are compositionally and transcriptionally distinct

The prolonged residence of developing B cells within discrete dura mater niches suggested the presence of a specialized cellular network capable of supporting local B cell development—potentially analogous to the stromal niches governing B lymphopoiesis in the bone marrow^19,58–62^. To gain a comprehensive view of the cellular and molecular interactions underpinning dural B lymphopoiesis—and to directly compare these to those present in early-life bone marrow niches—we analyzed non-immune cells from our P14 scRNA-seq dataset (**Figure S3B**). Across dural, calvarial bone marrow, and tibial bone marrow compartments combined, we identified populations of mesenchymal stem and progenitor cells (MSPC, *Lepr^high^, Adipoq^+^*), osteolineage cells (OLC, *Sp7^+^Mmp13^+^*), fibroblast-like cells (FLCs, *Pdpn^+^Foxp2^+^*), mural cells (*Rgs5*^+^*Mcam^+^*), and endothelial cells (*Pecam1*^+^*Flt1*^+^), among other populations that compose putative hematopoietic niches in these organs (**Figure 5A, Figure S6A**). Endothelial cells, MSPCs, and OLCs—which serve as niche cells supporting multilineage hematopoiesis—were abundant in bone marrow non-immune compartments as previously described^19,61,63,64^ (**Figure 5B-C**). However, the P14 dura contained few cells that resembled canonical MSPCs/OLCs; instead, FLCs and mural cells predominated the dural stromal landscape. Intriguingly, we found that sinus-proximal dural B cell clusters at P14 were enmeshed in dense networks of ERTR7^+^ fibers, likely representing FLCs and the collagen components they produce^65^ (**Figure 5D**).

**Figure 5.**
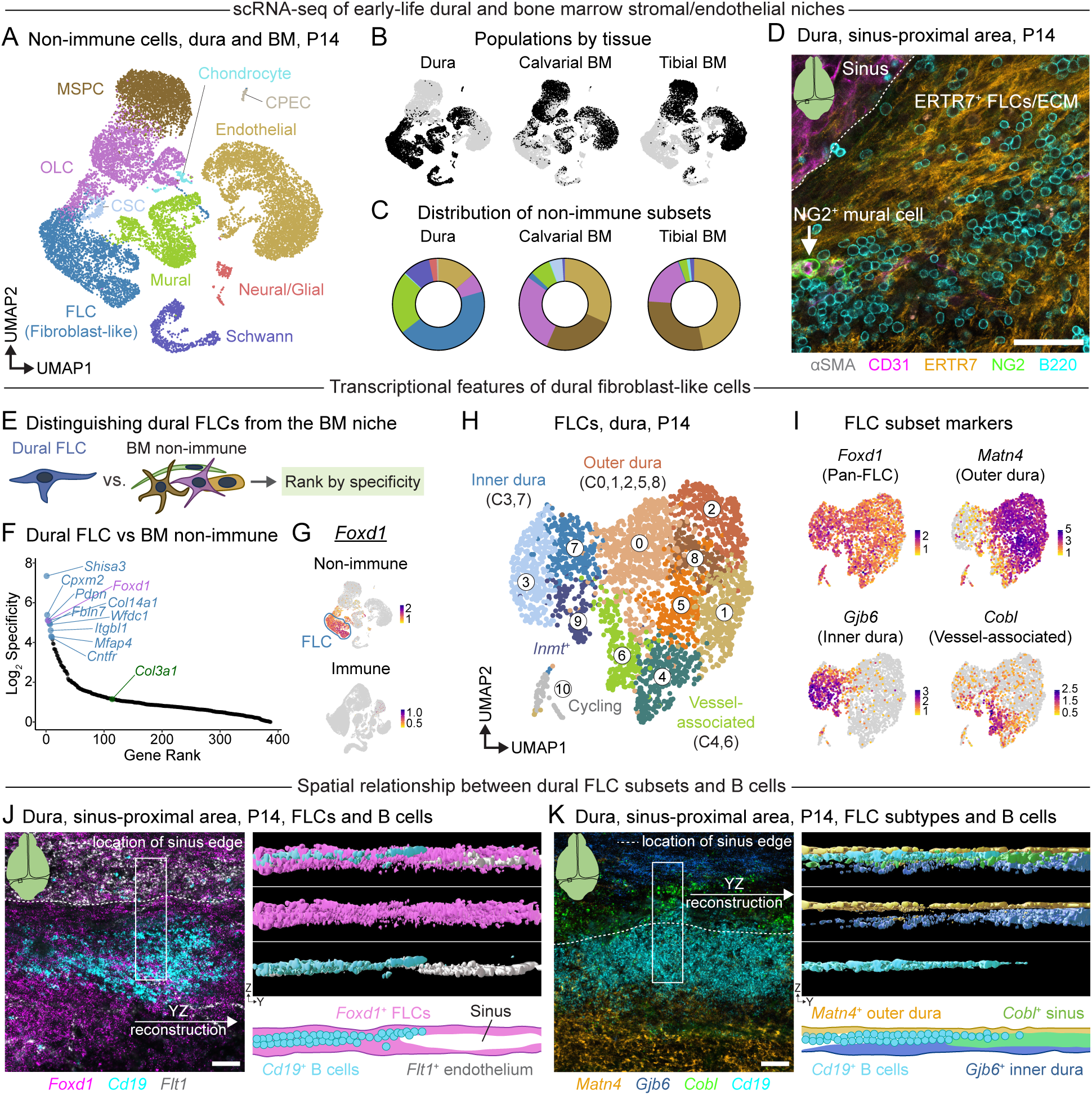
Anatomically and molecularly distinctive fibroblast-like cells compose the dural B cell niche. (A) UMAP depicting scRNA-seq data of dural, calvarial bone marrow (BM), and tibial bone marrow non-immune cells at P14. n = 3 mice from data presented in Figure S3. MSPC—Mesenchymal stem and progenitor cell; OLC— Osteolineage cell; FLC—Fibroblast-like cell; Mural cell (smooth muscle cell and pericyte); Chondrocyte; CSC— Cranial stromal cell; Endothelial cell; Schwann cell; Neural/glial cell (pinealocyte, astrocyte-like cell, neuronal cell); CPEC—Choroid plexus epithelial cell. (B) UMAPs split by tissue corresponding to (A). (C) Distribution of non-immune cell subsets per tissue. (D) Representative image of B cell association with stromal and endothelial cell types in the transverse sinus-proximal area at P14. αSMA—α-smooth muscle actin, white; CD31—vasculature, magenta; ERTR7—FLCs/extracellular matrix (ECM), orange; NG2—neural/glial antigen 2, mural cells, green; B220—B cells, cyan. Scale bar represents 50 µm. (E) Identification of genes enriched in and specific for dural FLCs compared to bone marrow non-immune cells. Differentially expressed genes upregulated in dural FLCs were assigned a specificity score where Specificity = % expression in FLCs/% expression in bone marrow non-immune cells. This list was further filtered for genes expressed in at least 70% of dural FLCs to identify pan-FLC markers that were specific for dural FLCs. (F) Dural FLC-enriched genes ranked by specificity and plotted on a log_2_ scale. The top 10 genes specific to dural FLCs are labeled (top 10, blue; gene of interest *Foxd1*, purple), as well as the global fibroblast marker *Col3a1* (green). (G) Expression of dural FLC-enriched gene *Foxd1* in dura and bone marrow non-immune and immune cells. FLCs are highlighted with a lasso. (H) UMAP of dural FLCs at P14. Individual clusters (e.g. C0, C1, etc.) and broad FLC groupings corresponding to anatomical divisions in the dura (outer dura, inner dura, vessel-associated) are indicated. (I) Expression of pan-dural FLC gene, *Foxd1*, and FLC subtype markers *Matn4*, *Gjb6*, and *Cobl*. (J) Association of *Cd19*^+^ B cells with *Foxd1*^+^ FLCs in the transverse sinus-proximal area at P14. mRNA transcripts were labeled using RNAscope. Image represents a maximum intensity projection of 3 µm from a confocal Z stack (left). Scale bar represents 50 µm. The full Z stack was used to reconstruct a portion this region in 3D in Imaris. Surfaces were constructed out of each channel and then used to visualize the location of B cells relative to FLCs and the sinus in the YZ dimension (right). A diagram depicting the approximate location of B cells in relationship to FLCs and the transverse sinus is also shown. *Foxd1*—FLCs, magenta; *Cd19*—B cells, cyan; *Flt1*—endothelial cells, white. (K) Association of *Cd19*^+^ B cells with dural FLC subsets composing the outer dura (*Matn4*), inner dura (*Gjb6*), and sinus/vessel-associated dura (*Cobl*) in the transverse sinus-proximal area at P14. Image represents a maximum intensity projection of 3 µm from a confocal Z stack localized to a B cell cluster near the transverse sinus (left). Scale bar represents 50 µm. The full Z stack was used to reconstruct a portion this region in 3D in Imaris. Surfaces were constructed out of each channel and then used to visualize the location of B cells relative to FLC subtypes in the YZ dimension (right). A diagram depicting approximate location of B cells and dural FLC subtypes is also shown. Note that there is some *Cobl* expression outside of the sinus and below the B cell cluster that is not depicted in the diagram. *Matn4*—outer dura FLCs, orange; *Gjb6*—inner dura FLCs, blue; *Cobl*—sinus-associated FLCs, green; *Cd19*—B cells, cyan.

Given this intimate association between FLCs and B cells, we next investigated the putative function of dural FLCs as components of the niche supporting early-life B cell development. To this end, we leveraged our scRNA-seq data to identify unique transcriptional features of dural FLCs that would allow us to genetically target them and interrogate their role in early-life B lymphopoiesis independently of medullary niches. We calculated differentially expressed genes between dural FLCs and other non-immune cells (e.g. bone marrow non-immune cells), filtered this list for genes expressed in at least 70% of dural FLCs, and ranked the filtered list by specificity for dural FLCs (**Figure 5E**). In these analyses, specificity represented the percentage of FLCs expressing a given gene divided by the percentage of other cells expressing the gene. This approach identified dural FLC-enriched genes—including *Shisa3* and *Foxd1*—that ranked highly for FLC specificity against non-immune cells in the bone marrow (**Figure 5F**) and against other non-immune cells in the dura (**Figure S6B-C**). This contrasted with other canonical fibroblast genes such as *Col3a1*, which ranked lowly for specificity in dural FLCs despite high expression levels. The transcription factor forkhead box D1 (*Foxd1*) emerged as a top candidate for targeting dural FLCs, as it was mostly absent from other non-immune and immune cell populations in the dura and bone marrow (**Figure 5G**). We also mined publicly-available scRNA-seq data describing fibroblasts/stromal cells in multiple murine organs^66^, revealing that *Foxd1* was largely absent from other major lymphoid organ and non-lymphoid, non-CNS, organ fibroblast/stromal cell populations across the body (**Figure S6D**). Consequently, we used a *Foxd1*^GFP-Cre^ mouse strain^67^ to target dural FLCs, revealing efficient recombination in these cells and stromal populations across dura and other brain border regions (**Figure S6E–H**). Brain border stromal cells showed elevated *Foxd1*^GFP-Cre^-mediated targeting compared to stromal and endothelial cells in the calvarial bone marrow, tibial bone marrow, and spleen, establishing the *Foxd1*^GFP-Cre^ strain as a reliable genetic tool for preferential manipulation of brain border FLC and stromal niches.

### Molecularly distinctive FLC subtypes compose the sinus-proximal B cell niche

Although several studies have described transcriptionally distinctive populations of dural FLCs that form the borders of this tissue (e.g. inner dura—or dural border—FLCs, and outer dura—or periosteal—FLCs) ^36,68–70^, little is known about the anatomical or molecular organization of FLCs in the sinus-proximal area of the early-life dura. To address this gap—and to determine whether specific subsets of FLCs associate with dural B cells—we further dissected the P14 dural FLC compartment using single cell sequencing and spatial approaches. Indeed, these analyses revealed several broad groups of FLCs that shared transcriptional features with previously described, geographically distinct, dural FLC populations (**Figure 5H**). While all FLC populations expressed *Foxd1, Shisa3,* and the canonical fibroblast marker *Col3a1*, outer dura/periosteal dura FLCs showed elevated expression of *Matn4*, *Comp, and Scara5* (**Figure 5I**, **Figure S6I**). Inner dura/dural border FLCs preferentially expressed transcripts encoding junctional proteins including *Gjb6*, *Gjb2,* and *Mpzl2*, as well as those encoding the solute transporters *Slc47a1* and *Slc5a6*. We also identified a group of FLCs enriched for select markers found in perivascular stromal cells and/or endothelial cells—including *Notch3*, *Col4a1*, and *Edn3*^71^—as well as markers specific to these putative vessel-associated FLCs compared to other dural stromal cells, such as *Cobl* and *Prss35* (**Figure 5I**, **Figure S6I**).

We next examined the localization of these dural FLC subtypes by RNAscope, aiming to elucidate the spatial relationship between distinct dural FLC subsets and B cells. This approach confirmed that dense networks of *Foxd1*^+^ FLCs enveloped B cell clusters in the sinus-proximal area of the dura (**Figure 5J**). Further, co-labeling for FLC subtype markers *Matn4, Gjb6, and Cobl* revealed an intricately organized FLC compartment at P14. *Cobl*-expressing FLCs lined the dural sinuses, whereas *Matn4*^+^ FLCs formed a distinct layer on the dorsal side of the dura and *Gjb6*^+^ FLCs resided mainly on the ventral side of the sinus-proximal dura (**Figure 5K**). Interestingly, *Cd19*^+^ B cells occupied a distinct physical space demarcated on the dorsal side by *Matn4*^+^ outer dura FLCs, the ventral side by *Gjb6*^+^ inner dura FLCs, and the lateral side by *Cobl*^+^ sinus-associated FLCs. Thus, the dural B cell niche comprises FLCs that can be distinguished from bone marrow niche cells by their expression of *Foxd1* and occupies a distinct anatomical location at the interface of previously described layers of the dura and the dural sinuses.

### A developmentally regulated dural FLC niche orchestrates B lymphopoiesis via CXCL12

To define the molecular interactions supporting early-life B cells in this unique dural niche, we performed ligand-receptor analysis between non-immune cells and B cells present in our P14 dura and bone marrow scRNA-seq datasets (**Figure S7A**). This approach allowed us to identify candidate interactions between dural FLCs and B cells, uncover additional communication pathways between B cells and other niche cells in the dura, and directly compare these putative interactions with those occurring between niche cells and B cells in the bone marrow. In the dura, we identified numerous non-immune cell–to–B cell interactions, many of which overlapped with niche–to–B cell interactions found in the bone marrow (**Figure S7B**). Indeed, dural and bone marrow non-immune cells interacted with B cells via ligands that are known to regulate B lymphopoiesis^72^ (e.g. *Cxcl12-Cxcr4, Vcam1-Itga4+Itgb1, and Kitl-Kit*) as well as other pathways including those between extracellular matrix (ECM) components and B cell adhesion molecules (e.g. *Col1a1-Sdc1* and *Fn1-Itga4+Itgb1*) (**Figure S7B**). Surprisingly, however, we did not recover an interaction between non-immune cell-derived *Il7* and B cell *Il7r* in the dura, whereas this interaction appeared in the calvarial and tibial bone marrow compartments in our dataset. As both CXCL12 and IL-7 critically regulate B lymphopoieisis^19,58,59,62,73–77^, we further investigated the cell types expressing these factors in the dura and bone marrow. This analysis revealed *Cxcl12* expression in several dural cell types including FLCs, mural cells, and endothelial cells, and in bone marrow MSPCs and OLCs as previously described^25,63,78,79^ (**Figure S7C**). The cognate receptor for *Cxcl12*, *Cxcr4*, appeared in both dural and bone marrow B cells. *Il7* was absent in dural cell types captured in this dataset but highly expressed in MSPCs and OLCs in the bone marrow (**Figure S7D**). In contrast, both dural and bone marrow B cells expressed its cognate receptor, *Il7r*.

We focused on the chemokine CXCL12 for further analyses as it showed prominent expression in dural FLCs and plays an established role in medullary B lymphopoiesis^19,58,59,62,73,74^. Indeed, examination of *Cxcl12*^dsRed2+^ mice revealed that FLCs composed nearly 80% of *Cxcl12*^dsRed2+^ stromal and endothelial cells in the dura at P14 (**Figure S7E-F**). To assess the spatial distribution of *Cxcl12*-expressing FLCs in the dura, we crossed *Foxd1*^GFP-Cre^ mice—which label dural FLCs—to *Cxcl12*^dsRed2^ mice and prepared dural whole mounts co-labeled for B cells and vasculature. Dense networks of *Foxd1*^GFP+^*Cxcl12*^dsRed2-high^ FLCs occupied the sinus-proximal area at P15, intercalating into B cell clusters and wrapping their processes around individual B cells (**Figure 6A**). Co-labeling of *Cxcl12* and FLC subtype markers *Matn4, Gjb6,* and *Cobl* allowed us to further evaluate the cellular and spatial distribution of this *Cxcl12*^+^ FLC niche. While FLCs in each anatomically defined compartment expressed some *Cxcl12*, a large proportion of *Cxcl12* signal filled the physical space enclosed between the inner dura, outer dura, and sinus-associated FLC compartments (**Figure S7G-H**). Indeed, *Cd19*^+^ B cell clusters co-occupied this distinct anatomical niche (**Figure 6B-C**), suggesting that *Cxcl12* may represent one mechanism by which FLCs coordinate B cell localization to the sinus-proximal area of the early-life dura.

**Figure 6.**
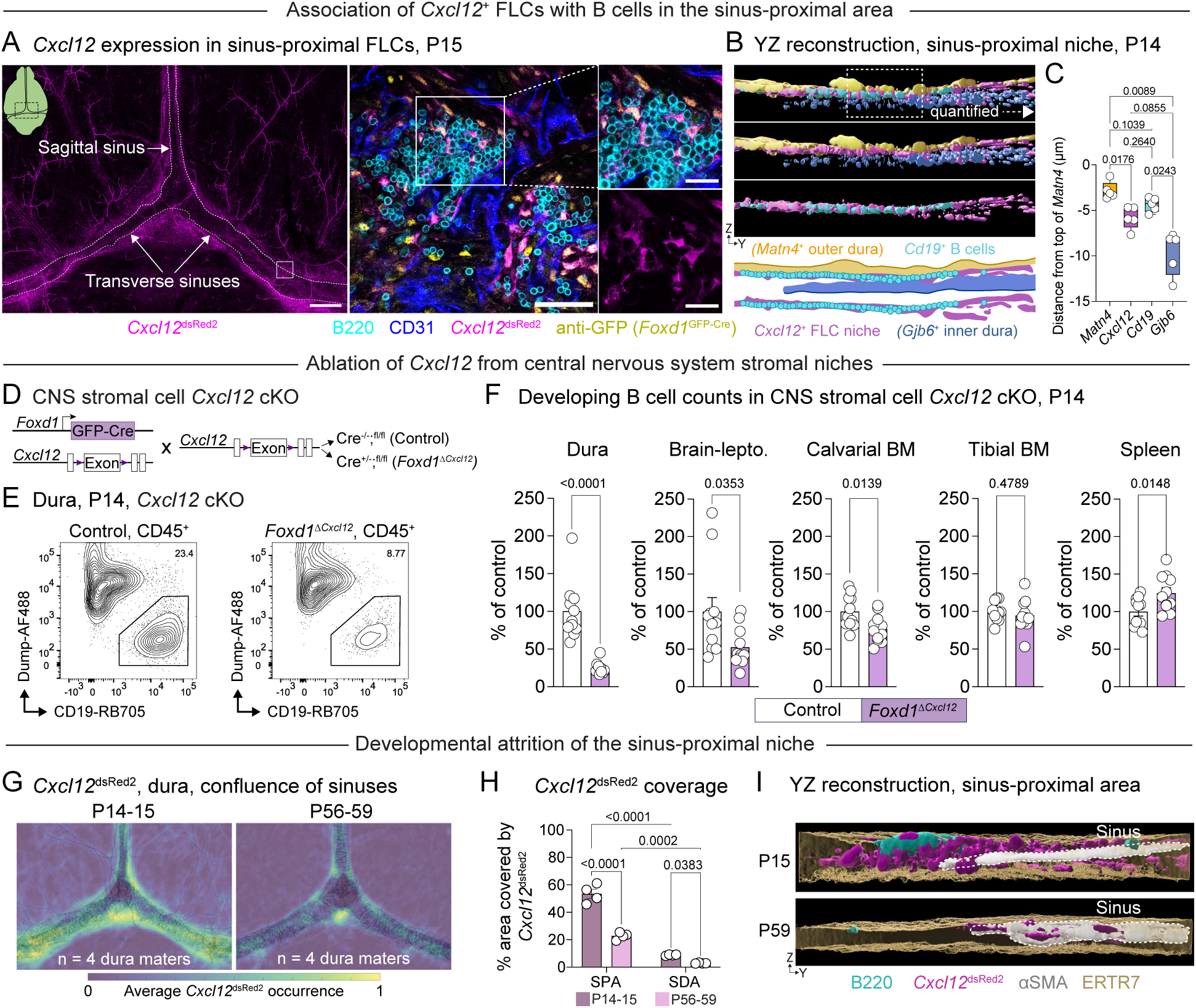
Dural fibroblast-like cells orchestrate local B lymphopoiesis via CXCL12. (A) Representative tile scan of a whole mount dura from a P15 *Foxd1*^GFP-Cre^;*Cxcl12*^dsRed2^ mouse depicting the confluence of the sinuses (left). High magnification image and cropped insets of a B cell cluster along the transverse sinus/bridging vein (right). B220—B cells, cyan; CD31—vasculature, blue; *Cxcl12*^dsRed2^—*Cxcl12*, magenta; anti-GFP staining of *Foxd1*-driven GFP, yellow. The white box on the tile scan indicates the approximate area depicted in the high magnification image. The tile scan represents a Z-projection of multiple stacks spanning the depth of the tissue, scale bar 500 µm. The high magnification images represent single plane confocal images, scale bar 50 µm for the high magnification image and 20 µm for the cropped insets. (B) Association of *Cd19*^+^ B cells with *Cxcl12* and dural FLC subsets composing the outer dura (*Matn4*) and inner dura (*Gjb6*) at P15. A series of Z-stack confocal images were taken of the sinus-proximal area corresponding to the location of a B cell cluster and then reconstructed in 3D in Imaris. Surfaces were constructed out of each channel and then used to visualize the location of B cells relative to *Cxcl12* and FLC subtypes in the YZ dimension. The boxed area indicates the ROI quantified in (C). A diagram depicting approximate location of B cells, the *Cxcl12*^+^ niche, and dural FLC subtypes is also shown with and without the inner/outer dura layers. *Matn4*—outer dura FLCs, orange; *Cxcl12*—niche signal, magenta; *Gjb6*—inner dura FLCs, blue; *Cd19*—B cells, cyan. (C) Quantification of the Z position of dural FLC layers, the *Cxcl12*^+^ niche, and B cells relative to the dorsal surface of the dura (demarcated by the highest *Matn4*^+^ point), related to Imaris reconstructions in (B). Each data point represents the relative median Z position of signal from a given channel at the center of a transverse sinus B cell cluster, averaged across two clusters from one mouse. n = 5 mice. Welch’s ANOVA with Dunnett’s T3 multiple comparisons test. F (3.000, 8.534) = 12.99, P = 0.0015. (D) Conditional knockout (cKO) of *Cxcl12* in *Foxd1*-expressing CNS FLCs/stromal cells using a *Foxd1*^GFP-Cre^ mouse line. (E) Representative flow cytometry plot depicting CD19^+^ B cells in the dura of Control (*Cxcll2^ff^*) or *Foxdl*^Δ^*^Cxcl12^* (*Foxd1*^GFP-Cre/+^;*Cxcl12*^fl/fl^) mouse at P14. Population is gated on live, single, extravascular, CD45^+^ cells. (F) Flow cytometric quantification of developing (CD19^+^CD20^-^IgM^-^IgD^-^) B cells across organs at P14 in Control (*Cxcll2*^fl/fl^) and *Foxdl^ΔCxcl12^* (*Foxdl*^GFP-Cre/+^;*Cxcll2*^fl/fl^) conditional knockout (cKO) mice. n = 11 control and 11 *Foxdl^Δ Cxcl12^* mice pooled from two independent experiments each consisting of one litter containing Control and *Foxdl^Δ Cxcl12^* littermates. B cell counts for each organ were normalized to the mean count for the Control group in each experiment. Error bars indicate SEM. Welch’s t test, two-tailed. Brain-lepto.—pooled brain and leptomeninges; BM—bone marrow. (G) Heatmaps depicting spatial localization of *Cxcl12* signal in the dura of P14-15 or P56-59 *Cxcl12*^dsRed2^ mice. Images from P14-15 or P56-59 mice were merged to create one heatmap for each age that represents the average occurrence of *Cxcl12*^dsRed2^ signal at each given XY coordinate across the dura. Heatmaps are overlaid with a composite image of aSMA signal (black outline) from the same mice at each age. n = 4 mice per age. (H) Percentage of the sinus-proximal area (SPA) and sinus-distal area (SDA) that is occupied by *Cxcl12* signal in the dura of P14-15 or P56-59 *Cxcl12*^dsRed2^ mice. n = 4 mice per age from (G). Two-way repeated measures ANOVA with Uncorrected Fisher’s LSD. F (1, 6) = 51.84, P = 0.0004. (I) *Cxcl12*^dsRed2^ expression in the sinus-proximal area at P15 or P59, related to (G). A series of Z-stack confocal images were taken of the sinus-proximal area and then reconstructed in 3D in Imaris. Surfaces were constructed out of each channel and then used to visualize the location of B cells and *Cxcl12* relative to the sinus. B220—B cells, cyan; *Cxcl12*^dsRed2^—*Cxcl12*^+^ niche, magenta; αSMA—α-smooth muscle actin, white. A transparent surface depicting FLCs/ECM (ERTR7, gold) is also included to demarcate the boundaries of the tissue.

To test the hypothesis that FLC-derived CXCL12 functionally regulates early-life B cell development we crossed *Foxd1*^GFP-Cre^ mice to *Cxcl12*^flox^ mice, generating *Foxd1*^1*Cxcl*12^ mice that targeted deletion of *Cxcl12* to dural and other CNS-associated FLCs/stromal cells (**Figure 6D**). To validate this knockout, we performed BaseScope for *Cxcl12*, revealing a near complete loss of *Cxcl12* signal from the dura of *Foxd1*^1*Cxcl*12^ mice at P14 (**Figure S7I**). We next quantified developing B cell subsets in the dura, brain-leptomeninges, calvarial bone marrow, tibial bone marrow, and spleen by flow cytometry at P14. *Foxd1*^1*Cxcl*12^ mice showed a pronounced decrease in numbers of developing B cells in the dural and brain-leptomeningeal compartments compared to *Cxcl12*-sufficient littermate controls (**Figure 6E-F, Figure S7J**). This loss occurred in Pro B, Pre B, and CD20^+^ newly formed/mature B cell populations (**Figure S7K**). We also observed a modest decrease in B cell numbers in the calvarial bone marrow, likely resulting from partial targeting of this niche by *Foxd1*-driven Cre (**Figure S6H**). However, deletion of *Cxcl12* in *Foxd1*-expressing cells did not alter B cell numbers in the tibial bone marrow, and the spleen showed slightly elevated numbers of developing B cells in *Foxd1*^1*Cxcl*12^ mice compared to *Cxcl12*-sufficent controls (**Figure 6F, Figure S7J**). These results demonstrate that *Foxd1*-expressing FLCs/stromal cells critically support B lymphopoiesis in early-life CNS border tissues via CXCL12, functioning largely independently of other lymphopoietic niches across the body.

Finally, we asked whether developmentally regulated changes in the *Cxcl12*^+^ FLC niche may, in part, explain the temporal dynamics of early life dural B lymphopoiesis. Heatmapping and quantification of *Cxcl12* localization in *Cxcl12*^dsRed2^ mice at P14-15 and P56-59 revealed a substantial decrease in dural *Cxcl12* expression from early-life to adult ages (**Figure 6G-H**). This niche remodeling was particularly apparent in the sinus-proximal area, where the dense networks of *Cxcl12*^high^ FLCs present at P14 largely disappeared by adulthood (**Figure 6I**). Taken together, these data suggest that sinus-proximal FLCs are exquisitely tuned to support B cells during the early-life window, producing the pro-hematopoietic factor *Cxcl12* during a discrete postnatal period to orchestrate this early-life wave of B lymphopoiesis.

## DISCUSSION

Here, we provide a detailed temporal map of brain border immune cell development and discover that the dura functions as a transient niche for B lymphopoiesis in the early-life window. We show that developing B cells populate the dura as a wave spanning the first postnatal month, participating in a coordinated, multi-organ wave of extramedullary lymphopoiesis that establishes the nascent mature B cell compartment. To define the niches that orchestrate dural B cell development, we profile early-life dural stromal cells and compare these to known populations of stromal cells that support B lymphopoiesis in the bone marrow. We identify molecularly distinctive populations of FLCs that can be distinguished from other early-life stroma by their expression of *Foxd1* and form anatomically defined niches for B cells in the sinus-proximal area of the dura. Sinus-proximal FLCs show elevated expression of the pro-hematopoietic chemokine *Cxcl12* in the early-life window, and ablation of *Cxcl12* from *Foxd1*-lineage stromal cells severely impairs local B lymphopoiesis in brain border tissues. Together, these data identify a critical function for dural FLCs in orchestrating early-life B cell development and provide a framework for studying how extramedullary stromal niches shape immune cell development across the body in the early-life window.

### Extramedullary organs as niches for early-life B lymphopoiesis

In mammals, the sites of B cell generation change across the lifespan, bookended by the fetal liver in embryonic development and the bone marrow in adulthood^5^. Our findings add to emerging evidence in both mice and humans that early-life extramedullary organs function as niches for B lymphopoiesis during the transition between fetal and adult blood cell production. For example, developing B cells populate the murine spleen and small intestine lamina propria during the first weeks of postnatal development^12,50^, and B cell progenitors appear in the human gut, skin, kidney, thymus, lymph nodes, spleen, and meninges during the first and second trimesters^13,80^. Our data further demonstrate that developing B cells populate CNS border tissues as a distinct early-life wave (Figures 2-3), originating from a common pool of embryonic progenitors that generate B cells in multiple early-life organs (Figure 4). In the dura, these progenitors expand extensively *in situ* to form discrete lymphopoietic hubs, suggesting that this niche can support prolonged, local B cell development. Thus, we posit that brain border tissues and other extramedullary organs do not merely serve as rest-stops for migrating B cell progenitors but may actively contribute to B cell output in early life.

In adulthood, immature B cells typically exit the bone marrow and transit to the spleen where they differentiate through transitional B cell stages to generate mature B cells^72^. We demonstrate that B cells born during the early life lymphopoietic wave contribute acutely to follicular, marginal zone, and to a lesser, extent B1a B cell compartments, and that B cells born after this wave no longer generate marginal zone B cells (Figure 3). These results dovetail with recent work demonstrating that B cells pulse-labeled between P10 and P19 persist in the adult splenic marginal zone B cell, B1a B cell, and lamina propria IgA^+^ plasma cell compartments, whereas early-life follicular B cells are largely replaced by adulthood^81^. Thus, B cells born during the early postnatal window likely contribute acutely and rapidly to the generation of mature B cell compartments that are then turned over at different rates. The observations that marginal zone B cells labeled in the early postnatal window persist into adulthood^81^ and are not appreciably generated after this early life lymphopoietic wave (Figure 3) suggest that early postnatal life is a critical window during which the composition and function of this compartment are shaped.

How B cells born during this early-life lymphopoietic wave may subsequently contribute to early-life antibody responses is unknown. Several studies have described formation of B cell-rich spontaneous lymphoid structures in early-life barrier tissues including the gut and lung in mice and humans^16,43,44^. It is possible that B cells in these structures derive directly from this early life lymphopoietic wave in anticipation of environmental exposure to pathogens. While we did not observe tertiary lymphoid-like structures in the early postnatal dura (Figure S2), examination of mice with a more diverse history of microbial experience^82^ or those infected with CNS-tropic pathogens^83,84^ could advance our understanding of how the CNS B cell compartment adapts to meet challenges commonly faced in the early-life window.

### Divergent mechanisms underpinning early-life and adult B lymphopoiesis in the dura

Recent work from several groups suggests that the adult dura harbors populations of developing B cells^31–33^. While we do observe a small number of developing B cells in the adult dura, our data suggests that most CNS-born B cells are generated in the first few weeks of postnatal life (Figure 3). B lymphopoiesis in the early-postnatal versus adult dura may indeed represent two distinct phenomena regulated by differential input from B cell progenitors. We show that embryonic progenitors coordinately seed the dura and other early-life organs to initiate early-life B lymphopoiesis and subsequently proliferate locally in distinct lymphopoietic foci to generate dural B cells (Figure 4). In contrast, developing B cells in the adult dura are independent of circulating cells^31–33^ and instead potentially derive from the calvarial bone marrow^31^. Neither our data nor other published work notes pronounced lymphopoietic foci in the adult dura. Instead, B cells in the adult dura are sparsely distributed along the dural sinuses, suggesting sporadic migration of developing B cells to these locations rather than prolonged development in defined microanatomical niches. Whether developing B cell foci re-emerge in adult dural niches during inflammatory events that induce extramedullary lymphopoiesis^85^ represents an interesting future question.

### Fibroblast-like cells as B cell and immune cell niches in CNS border tissues

In adult mammals, specialized populations of bone marrow mesenchymal stromal and osteolineage cells support nascent B cells^19,58,59,62^. We demonstrate that early-life dural B cells associate with FLCs that are transcriptionally distinct from bone marrow stromal cell populations and that occupy a unique anatomical niche bounded by the inner dura, outer dura, and dural sinuses (the sinus-proximal niche) (Figure 5). Building on previous studies^36,68–70,80,86^, we describe the transcriptional and spatial organization of FLC sub-populations that compose these distinct sinus-proximal dural compartments. Further, we identify genes—including *Foxd1*—that distinguish dural FLCs from bone marrow stromal, lymphoid organ stromal, and other organ specific FLC populations, allowing functional manipulation of the dural FLC/stromal niche using a *Foxd1*^GFP-Cre^ line (Figure S6). In general, the abluminal space of the dural sinuses serves as a niche for immune cells across the lifespan^25,27,30,34,35,41^, but we know little about the functional contributions of distinct stromal cell populations to immune regulation there. For example, how outer dura, inner dura, and vessel-associated FLC populations are remodeled—and whether distinct subpopulations contribute to maintenance of different immune cell populations—during development, after infection, or in the face of neurodegenerative disease is unknown. The insight into the dural FLC niche provided here will inform efforts to study and manipulate FLC–immune cell interactions in myriad contexts.

Similarly to MSPCs and OLCs that regulate multi-lineage hematopoiesis in the bone marrow^58,59,61–64^, early-life sinus-proximal dural FLCs produce high levels of the chemokine *Cxcl12* (Figure 6). To test whether dural FLC-derived CXCL12 functionally regulates early-life lymphopoiesis, we genetically ablated *Cxcl12* from CNS stromal cells using a *Foxd1*^GFP-Cre^ mouse line. This manipulation led to pronounced impairment of B lymphopoiesis in the dura and other CNS border tissues while largely sparing B cell development in other organs (Figure 6). Overall, these results suggest that FLC/stromal cell populations in CNS border tissues critically regulate local B lymphopoiesis, further supporting the idea that this process proceeds largely independently of B lymphopoiesis in other early-life organs after initial seeding by a common pool of progenitors. Given the critical role of CXCL12 in maintaining multiple stages of hematopoietic progenitors and early immune cells in the bone marrow^19^, we speculate that deletion of CXCL12 in CNS FLCs/stromal cells impairs CNS B lymphopoiesis through direct effects on B cells and by impacting initial localization of hematopoietic progenitors to the sinus-proximal niche. The effect of this deletion on recruitment of other immune cells to the early-life dura in development and disease—and the overall functional impact of this impairment—present intriguing areas for future investigation.

Additionally, the cellular sources of other factors that regulate B lymphopoiesis in the dura remain unexplored. For example, neither early-life dural stromal cells nor other dural cell types we examined express the early B cell growth factor *Il7* (Figure S7)—a cytokine critical for postnatal B cell development^75,77^. In contrast, early-life bone marrow MSPCs—and to some extent OLCs—produce high levels of IL-7 that likely work together with CXCL12 from the same cell types to coordinately regulate B lymphopoiesis^58,61,63,64^. We speculate that early-life dural IL-7 either derives from another local source that we did not examine—for example, lymphatic endothelial cells, which produce IL-7 in other organs^87^—or from an extra-dural source such as the blood or the calvarial bone marrow, which contains abundant IL-7–producing MSPCs (Figure S7). Further elucidating these and other interactions guiding B cell development in the early-life dura will inform our understanding of how extramedullary niches regulate B lymphopoiesis in the absence of specialized stromal cells found in primary lymphoid organs.

Collectively, our findings uncover a transient wave of B lymphopoiesis in the early postnatal dura and define the stromal niche that functionally supports it. In doing so, this study redefines developing brain border tissues as potential contributors to the establishment of the systemic adaptive immune system. It also underscores the broader principle that extramedullary stromal niches support early-life B lymphopoiesis and shape immune cell development across the body.

### Limitations of this study

Although we demonstrate that B cells born during the early life lymphopoietic wave contribute to the mature B cell compartment, we cannot trace or manipulate B cells derived from individual lymphopoietic tissue niches. For example, it would be informative to determine the geographic fate of dura-born B cells as the lymphopoietic wave subsides, whether they persist into adulthood, and how they respond to subsequent infectious or autoimmune insults to the CNS. Additionally, tools to broadly label and/or ablate B cells born in medullary versus extramedullary organs would allow us to untangle the relative contributions of these lymphopoietic niches to the overall function of the peripheral B cell pool. Although we propose that dural FLC-derived CXCL12 acts on early-life B cells to regulate their local development, we cannot exclude the possibility that constitutive ablation of CXCL12 from CNS stromal cells also alters other important aspects of dural niche development that have secondary effects on B cell development. For example, CXCL12 regulates embryonic blood and lymphatic vascular patterning^88–90^, which could indirectly affect the physical or molecular architecture of early-life dural niches. Tools designed to allow temporal control of gene expression in dural FLCs will help to address this limitation. Finally, this study is limited to experiments conducted in mice; whether the dura functions as an early-life niche for B lymphopoiesis in other species including humans is a question that merits future exploration.

## METHODS

### Mice

All experiments were conducted under protocols approved by the Boston Children’s Hospital Institutional Animal Care and Use Committee adhering to NIH guidelines for the humane treatment of animals. Mice were housed under a 12-hour light:12-hour dark cycle in Optimice cages (Animal Care Systems, C79112PF). *Ad libitium* access to food and water was provided, and environmental conditions maintained in accordance with guidance from the Association for Assessment and Accreditation of Laboratory Animal Care. Transgenic mouse strains were either ordered from Jackson Laboratories or obtained from collaborators and then bred in house for use in experiments. C57BL/6J mice were bred in house or ordered from Jackson Laboratories and allowed to acclimate before use in experiments. C57BL/6J mice (JAX #000664) were used for experiments aimed at characterizing the early-life B cell wave and the dural stromal niche. To visualize B cells in whole mount dural preparations, we generated Cd19^Cre^;^lsl^tdTomato mice by crossing strains B6.129P2(C)-Cd19tm1(cre)Cgn/J (JAX #006785) and B6.Cg-Gt(ROSA)26 Sortm14(CAG-tdTomato)Hze/J (JAX #007914). *Rag1*^RFP-CreERT2^ mice (Tg(Rag1-RFP,-cre/ERT2)33Narl) were a generous gift from Dr. Chrysothemis Brown at Memorial Sloan Kettering Cancer Center and were originally generated by the National Laboratory Animal Center. This strain was used to visualize *Rag1*^+^ B cells by immunohistochemistry and crossed to ^lox-stop-lox^YFP mice (B6.Cg-Gt(ROSA)26Sortm3(CAG-EYFP)Hze/J (JAX #007903)) to allow fate mapping of developing B cells. CARLIN mice^54^ (B6;129S4-Col1a1tm4Fcam/Mmjax (JAX #067061)) were a generous gift from Dr. Fernando Camargo at Boston Children’s Hospital. These were crossed to tetracycline-inducible Cas9 mice (B6;129S4-*Gt(ROSA)26Sortm1(rtTA*M2)Jae Col1a1tm1(tetO-cas9)Sho*/J (JAX# 029415)) to generate CARLIN;M2-rtTA;TetO-Cas9 mice, allowing DNA barcode-based lineage tracing of early-life B cells. *Flt3*^EGFP-CreER^ mice were also obtained from Dr. Fernando Camargo at Boston Children’s Hospital and are described in Patel et al^91^. This strain was bred to ^lox-stop-lox^tdTomato mice (listed above, JAX #007914) to allow fate mapping of *Flt3*-expressing hematopoietic progenitors. *Flt3*^EGFP-CreER^ mice were also crossed to ^lox-stop-lox^Confetti mice (B6.129P2-Gt(ROSA)26Sortm1(CAG-Brainbow2.1)Cle/J (JAX #017492)) to allow assessment of *in situ* expansion of *Flt3*^+^ progenitor clones and early B cells. *Foxd1*^GFP-Cre^ mice (B6;129S4-Foxd1tm1(GFP/cre)Amc/J (JAX #012463)) were used to target CNS stromal cells. To assess Cre targeting, this strain was bred to ^lox-stop-lox^YFP mice (B6.129X1-Gt(ROSA)26Sortm1(EYFP)Cos/J (JAX #006148)). To visualize cells co-expressing *Foxd1* and *Cxcl12*, *Foxd1*^GFP-Cre^ mice were bred to *Cxcl12*^dsRed2^ mice (Cxcl12tm2.1Sjm/J (JAX #022458)). *Cxcl12*^dsRed2^ mice were also used on their own to visualize *Cxcl12*-expressing cells. To ablate *Cxcl12* in *Foxd1*-expressing CNS stromal cells, *Foxd1*^GFP-Cre^ mice were crossed to *Cxcl12*^flox^ mice (B6(FVB)-Cxcl12tm1.1Link/J (JAX #021773)). Transgenic mice were generally genotyped using services from Transnetyx. CARLIN mice were genotyped in house using previously-described primers^54^. In some cases, mice expressing fluorescent proteins (e.g. *Cxcl12*^dsRed2^ and *Rag1*^RFP-CreERT2^) were genotyped by examining relevant organs under a fluorescent dissection scope. For experiments involving C57BL/6J mice, numbers of male and female mice per group were matched where possible. For the scRNA-seq experiment, only male mice were used. For experiments involving transgenic mice, males and females were used as available in each litter.

### Timed pregnancies and induction for fate mapping and lineage tracing experiments

For both CreER-based fate mapping and CARLIN lineage tracing experiments requiring embryonic induction, timed matings were set in the evening and separated the following morning, which we designated as (E)0.5. For CreER-based models, recombination was induced by injection of Tamoxifen (Sigma-Aldrich, T5648) dissolved in corn oil (Sigma-Aldrich, C8267-500ML) or 4-hydroxytamoxifen (Sigma-Aldrich, H7904) dissolved in 100% EtOH (Fisher Scientific, 18-602-023) and further diluted in corn oil. For fate mapping experiments involving *Flt3*^EGFP-CreER;lsl^tdTomato mice, postnatal Cre recombination (P3, P7) was induced by an intraperitoneal injection of Tamoxifen at a dose of 50 mg/kg. When tamoxifen (50 mg/kg) was administered to pregnant dams at E17.5, Progesterone (Sigma-Aldrich, P3972) was co-administered at a dose of 25 mg/kg. For experiments involving *Flt3*^EGFP-CreER;lsl^Confetti mice, Tamoxifen and Progesterone were administered to pregnant dams at E17.5 as a single dose of 100 mg/kg and 50 mg/kg, respectively. *Flt3*^EGFP-CreER;lsl^Confetti pups were delivered by cesarean section at E18.5 and fostered to a dam with an appropriately timed litter. For experiments involving *Rag1*^RFP-CreERT2^;^lsl^YFP mice, Cre recombination was induced in pups or juvenile mice by a single intraperitoneal injection of 4-hydroxytamoxifen at a dose of 50 mg/kg. For lineage tracing experiments using CARLIN;M2-rtTA;TetO-Cas9 mice, barcoding was induced by a single injection of doxycycline hyclate (Sigma-Aldrich, D9891) dissolved in 0.9% saline (Teknova, S5825) at a dose of 50 mg/kg. Pregnant dams (iE12.5 and iE17.5) were injected retro-orbitally. P3 pups were injected intragastrically/intraperitoneally.

### Immunohistochemistry on dural whole mount preparations

Fluorescent immunohistochemical staining was performed on whole mount dural preparations to visualize the spatial relationship between B cells, other immune cells, and stromal cells. Mice were first anesthetized by intraperitoneal injection of Avertin (2,2,2-Tribromoethanl (Millipore Sigma: Cat# T48402) in 2-Methyl-2-butanol (Millipore Sigma; Cat# 240486) and Hank’s balanced salt solution (HBSS, Life Technologies, 14175-145) at a dose of approximately 800 μg/g body weight. In cases where vasculature was labeled intravenously, anesthetized mice received an intracardiac or retro-orbital injection of anti-CD31-AF647 (clone MEC13.3, 0.4 μg/g body weight, Biolegend 102516) or anti-CD31-BV421 (clone 390, 0.4 μg/g body weight, Biolegend 102424). Three minutes later, mice were transcardially perfused with at least 1 mL per gram body weight of ice cold HBSS. Mice were decapitated and skull caps were removed and drop fixed overnight in a formaldehyde-based fixative. Different fixation conditions were used depending on the intended staining panel for the experiment, including: 1:1 BD ICC Fixation buffer in HBSS (BD Biosciences, 550010) (approximately 1% final formaldehyde content); eBioscience IC Fixation buffer (eBioscience, 00-8222-49) (4% final paraformaldehyde, PFA, content); or 4% PFA (16% PFA, EM grade, Electron Microscopy Science, 15710) diluted in HBSS. Fixed tissues were washed in HBSS and stored at 4°C until further use. Dura maters were carefully dissected from skull caps and placed in a 24 well pate for staining. To permeabilize and block the tissues, dura maters were incubated in 300 μL of 0.3% Triton-X100 (Sigma-Aldrich, X100-100ML), 20% horse serum (Vector Laboratories S-2000-20) or donkey serum (Sigma-Aldrich, D9663-10ML), anti-CD16/CD32 (clone 2.4G2, 1:100, BD Biosciences, 553141), and HBSS for one hour at room temperature. After removal of the blocking solution, 300 μL of a primary antibody staining solution (primary antibodies of interest, Brilliant Stain Buffer Plus (BD Biosciences, 563794), 0.3% Triton-X100, 20% serum, and HBSS) was added and allowed to stain overnight at 4°C. Antibodies used are listed in **Table S1**. The following day, tissues underwent three wash steps for at least one hour each in 0.3% Triton-X100 in HBSS under slow agitation. For some panels, a secondary antibody cocktail diluted in 0.3% Triton-X100, 20% serum, and HBSS was applied, stained overnight at 4°C, and washed again the following day. Washed tissues were either then stained with DAPI diluted 1:10000 in HBSS or proceeded directly to tissue mounting. Dura maters were positioned on microscopy slides (Leica, 3800200) and smoothed out with paintbrushes until the tissues were almost dry. At this point, ProLong Diamond Antifade Mountant (Thermo Fisher, P36970) was added, and slides were cover slipped before imaging.

### RNAscope and Basescope on whole mount dural preparations

*In situ* hybridization using RNAscope technology (Advanced Cell Diagnostics, ACD) was employed to visualize B cells and stromal cell subsets in whole mount dural preparations. BaseScope technology (ACD) was used to evaluate the expression of *Cxcl12* exon 2 in dura from *Foxd1*^GFP-Cre^;*Cxcl12*^flox^ mice where deletion of that exon is targeted to CNS stromal cells. Skull caps were collected from mice transcardially perfused with ice cold HBSS and drop fixed for 24 hours in 4% PFA diluted in HBSS. Tissues were washed in HBSS and dura maters were carefully dissected from the skull caps. Dura maters were then flattened onto Superfrost Plus microscope slides (Fisher Scientific, 22-037-246) with the dorsal side facing up. Slides were dried for 15 minutes at 60°C on a slide warmer, frozen on dry ice, and stored at −80°C.

The RNAscope Multiplex Fluorescent V2 Assay was adapted for use on whole mount dural preparations, requiring substantial modifications to the original protocol to allow better target probe penetration and to reduce non-specific probe binding in the thick areas near the dural sinuses. Prepared slides were dried for one hour at 60°C and then rehydrated by immersing in HBSS at room temperature. The tissue then underwent additional fixation for one hour at room temperature in 4% PFA diluted in HBSS and was then permeabilized by immersing the slides in 0.3% Triton X-100 diluted in HBSS (HBSS-T) for one hour at room temperature. As an additional permeabilization step, the tissue underwent a dehydration series in increasing concentrations of ethanol diluted in HBSS-T (3:7, 1:1, 7:3), followed by immersion in 100% ethanol for 5 minutes at room temperature, and then rehydration by immersion in decreasing concentrations of ethanol diluted in HBSS-T (7:3, 1:1, 3:7) (adapted from Zimmerman et al.^92^). Dura maters were then treated with hydrogen peroxide for 5 minutes following manufacturer’s instructions, followed by Target Retrieval for 15 minutes. Tissues were then dehydrated by immersing the slides in 100% ethanol for five minutes at room temperature, followed by application of a hydrophobic barrier (ImmEdge hydrophobic barrier pen). The tissue was then incubated with Protease IV provided in the RNAscope Multiplex Fluorescent V2 Assay kit for two hours at room temperature to ensure optimal target probe accessibility. Protease-treated slides were washed in RNase free water for five minutes at room temperature before proceeding to hybridization.

Dura maters were hybridized with up to four target probes per experiment in the HybEZ Hybridization System at 40°C overnight. To ensure removal of unbound mRNA probes, hybridized slides were washed three times for 15 minutes in 1X RNAscope Wash Buffer. The tissue then underwent the amplification and signal development steps as suggested by ACD with extended wash steps (three times for 15 minutes each in 1X RNAscope Wash Buffer) after each fluorophore incubation step. Upon developing the signal for all target probes, Prolong Diamond Antifade Mountant was applied, and slides were cover slipped for imaging.

These modifications in tissue pretreatment, hybridization and washes for whole mount mouse dural preparations were also applied to the Basescope Assay. Relevant reagents for both protocols are listed in **Table S2**.

### Confocal and slide scanner imaging

Mounted tissues were imaged on a Zeiss LSM880 confocal microscope with Zen (black edition) 2.3 software or a Zeiss Axioscan 7 slide scanner with Zen 3.4 slide scanner software. Confocal images were acquired with up to four imaging tracks comprising seven distinct fluorescent channels. Lasers allowed excitation at 405 nm, 458 nm, 488 nm, 514 nm, 561 nm, 594 nm, 633 nm, and 730 nm. Fluorescent signal was collected by three internal detectors with adjustable band pass filters and two external detectors with fixed band pass filters in the far red and infrared range. Digital bandpass filter settings were optimized to allow resolution of the five channels collected on the internal detectors. Images were taken with 10x or 20x air objectives, or a 40x oil immersion objective, with pinhole aperture settings matched across channels for a given imaging paradigm. For quantified imaging data, imaging parameters including laser powers, detector gains, and pinhole apertures were kept the same for images within one batch. Images in single regions of interest were taken as confocal stacks covering the depth of the tissue or as single plane confocal images. Confocal tile scan overviews were acquired as Z stacks covering the depth of the tissue over an area of 7×5 tiles at 10x magnification centered around the confluence of the sinuses. These images were then stitched and converted into maximum intensity Z projections in Zen 2.3. The Zeiss Axioscan 7 slide scanner was used to generate overviews of dural whole mounts. Fluorescent or brightfield imaging modalities were used depending on the staining paradigm and images were taken using a 20x objective. Tile scans of entire dural whole mounts were generated using an automatic threshold-based tissue detection strategy. This approach selected the entire dura as the imaging region of interest, created a coarse focus map with a 5x objective using an “onion skin” focus map strategy, and imaged Z stacks for each tile that covered the depth of the tissue. The images were automatically stitched and converted into maximum intensity Z projections in Zen 3.4 slide scanner software.

### Image processing

Confocal images were further processed in Fiji 2.16.0^93^ or Imaris 10. Slide scanner images were further processed in Fiji. For confocal and slide scanner images processed in Fiji, display intensities were manually adjusted for each channel and then applied across a given batch of images. Images taken in single regions of interest are shown as single confocal planes or as a maximum Z projection of several contiguous Z planes (for example, covering a Z depth of 3 μm). Confocal and slide scanner tile scans are displayed as maximum intensity Z projections. Additionally, confocal tile scan images were used to generate heatmaps representing the location of B cells or *Cxcl12*-expressing cells in the dura (see “Quantification and heatmapping of B cells or *Cxcl12* in dura mater tile scan images”). Imaris was also used to reconstruct confocal Z-stack images in three dimensions. Minimum and maximum display ranges were adjusted per channel in each image. Then, a region of interest was chosen that encompassed either a B cell cluster or part of the sinus-proximal area of the dura. Surfaces were created based on the fluorescent signal from each channel in this region, which allowed visualization of the dural sinus and B cells, or dural FLC subsets and B cells, in three dimensions. In some cases, these surfaces were then used to calculate the median Z-position of different FLC layers and of B cells in Imaris.

### Quantification and heatmapping of B cells or *Cxcl12* in dura mater tile scan images

Tile scan images of dural whole mounts stained for αSMA (dural sinuses and arteries) and B220 (B cells) or *Cxcl12*^dsRed2^ (B cell niche component) were used to generate heatmaps of dural B cell or niche localization and to quantify B cell or niche signal coverage of different areas of the dura. The basis of this approach was recently described in collaborative work^40^ and has now been incorporated into a custom MATLAB^94^ graphic user interface (GUI) to allow user-friendly heatmapping and quantification of dural tile scan images with the use of the Image Processing Toolbox version 25.1 (R2025a)^95^. Maximum Z projected images were converted to TIFF format, imported to MATLAB version: 25.1.0 (R2025a), and split into individual fluorescent channels. αSMA images were then overlaid onto a reference vector image resembling the structure of the confluence of the dural sinuses and registered using MATLAB Control Point Selection Tool^95^ to assign moving and fixed points to the αSMA image and reference vector image respectively. Resultant registration coordinates were applied to the B220 or *Cxcl12*^dsRed2^ channel for each image, facilitating alignment of images from multiple mice based on the location of the dural sinuses. Autofluorescent signal from the pineal gland or technical aberrations in the images (e.g. bubbles) were then manually masked and removed from B220 and *Cxcl12*^dsRed2^ channels. All channels were then thresholded and binarized, applying a consistent threshold across all images in a batch for a given channel. Binarized images representing a given channel were aggregated across samples, creating a single image representing the arithmetic mean intensity per pixel for each experimental group. For B220 and *Cxcl12*^dsRed2^ channels, each aggregated image was then used to generate a pseudocolor plot, and interpolated shading was applied to each plot. Resulting heatmaps of the interpolated mean pixel intensity display the relative frequency with which B cell or *Cxcl12*^dsRed2^ signal occurs at a given location across dura maters from multiple mice within a given experimental group. These heatmaps were adjusted to 60% opacity in Adobe Illustrator (version 29.8.1) and overlaid with the corresponding aggregated αSMA image to visualize B cell or *Cxcl12*^dsRed2^ localization relative to the dural sinuses.

The registered and binarized αSMA channel was also used to segment the dura into two areas of interest (the sinus-proximal area and the sinus-distal area). The sinuses were manually traced using MATLAB drawfreehand()^95^ to create a mask covering the sinus. Once the sinus area was segmented, the mask was dilated by 300 μm from each edge of the sinus to generate a final mask of the sinus-proximal area. The sinus-distal area represents any area greater than 300 μm from the edges of the sinuses. Masks representing the pineal gland and imaging aberrations were then subtracted from the sinus-proximal area and sinus-distal area masks. By applying the final sinus-proximal and sinus-distal area masks to the registered and binarized B220 or *Cxcl12*^dsRed2^ channels, we calculated the total area of B220 or *Cxcl12*^dsRed2^ pixels contained in each region for each individual image. This metric was then used to calculate percent of total B cell signal contained in each region and the percent of each region occupied by B220 or *Cxcl12*^dsRed2^ signal (referred to as B cell or *Cxcl12*^dsRed2^ coverage).

### Quantification of *Cxcl12* BaseScope signal in dural whole mounts

Ablation of *Cxcl12* exon 2 in dural whole mounts from *Foxd1*^GFP-Cre^;*Cxcl12*^flox^ mice was visualized using BaseScope and quantified using QuPath^96^. Brightfield slide scanner images of whole mounted dura maters labeled for *Cxcl12* exon 2 using the BaseScope chromogenic assay were binarized in QuPath using an intensity-based thresholding for *Cxcl12* expression. A rectangular region of interest centered around the confluence of the dural sinuses was selected and the percent of this area covered by *Cxcl12* signal was quantified in *Foxd1*^GFP-Cre/+^;*Cxcl12*^fl/fl^ and *Cxcl12*^fl/fl^ animals.

### Preparation of tissues for flow cytometric analysis and cell sorting

One of several protocols was used to generate single cell suspensions for flow cytometry on CNS immune cells or stromal cells and corresponding populations in other organs, depending on the purpose of the experiment. These are listed below and encompass all flow cytometry experiments and the preparation of tissues for fluorescence activated cell sorting in CARLIN lineage tracing experiments. Tissue preparation for single cell RNA sequencing is discussed separately in “Preparation of tissues for single cell RNA sequencing.”

#### Intracardiac labeling of circulating immune cells

For most experiments, CD45^+^ leukocytes from the blood were labeled by intracardiac injection before transcardiac perfusion so that they could be later excluded during flow cytometric analysis. In anesthetized mice, the chest cavity was surgically opened to expose the heart and 0.2 μg/g body weight of anti-CD45-PE (clone 30-F11, Biolegend, 103106) was injected into the left ventricle and allowed to circulate for three minutes. A small amount of blood was then drawn from the right atrium and directly diluted into 10 ml of ice cold FACS buffer (0.5% bovine serum albumin (BSA, Sigma-Aldrich, A2153-1KG) and 2mM EDTA (Research Products International, E14000-100.0) diluted in HBSS).

#### Extraction of tissues

Mice were perfused with approximately 1 mL/g body weight of ice cold HBSS before tissue harvest. To access CNS tissue, perfused mice were decapitated and the skin of the head peeled forward toward the nose. Using spring scissors (Fine Science Tools, 91500-09), the skull was cut axially from the foramen magnum to the eye socket on each side and then between the eye sockets across the frontal bone. The skull cap was then removed and placed in ice cold RPMI-1640 (Thermo Fisher Scientific, 11835-030) + 10 mM HEPES buffer (Sigma Aldrich, H0887-100ML) (RPMI-H). The brain (including attached leptomeninges and choroid plexuses) was also removed at this point and stored in ice cold RPMI-H. Depending on the experiment, other organs including the spleen, tibia and femur bones, lymph nodes, small intestine, and liver were also collected for further processing.

#### Separation of the dura mater from the calvarial bone

To extract the dura mater, the skull cap was placed with the dura facing up in a dissecting dish with ice cold HBSS under a dissecting microscope. Spring scissors were used to cut axially around the skull cap in a line that goes directly under the auditory bulla, creating a consistent sized piece of calvarial bone and dura. To minimize contamination of the dura by cells from the calvarial bone marrow that are released when cutting through the bone, skull caps were then inverted, shaken several times, and transferred to a new dissecting dish containing fresh HBSS. Any remaining bone fragments were carefully removed from the interior aspect of the skull. The third ventricle choroid plexus (attached to the pineal gland) and the tentorium cerebelli were also removed when visible. The dura was then carefully peeled away from the skull and placed in a 1.5 mL microcentrifuge tube containing 500 μL of RPMI-H. In cases where the calvarial bone marrow was evaluated, the remaining skull cap was further processed by removing and discarding the occipital bone. The dorsal side of the skull cap was then cleaned of periosteum, and the skull cap was placed into a 2 mL microcentrifuge tube containing 750 μL of ice cold RPMI-H.

#### Removal of the epithelial layer from the small intestines

Small intestines were flushed with ice cold HBSS, opened longitudinally and rinsed in ice cold HBSS to remove residual luminal content, and incubated for 10 minutes in 4 mL of 1 mM Dithiothreitol (DTT) (Sigma-Aldrich, 10197777001) and 2% fetal bovine serum (FBS) (Invitrogen, 10437-028) in HBSS + 10 mM HEPES buffer (HBSS-H) on ice while periodically inverting. Intestines were then washed in ice cold HBSS and resuspended in 10 mL of 10 mM EDTA and 2% FBS in HBSS-H for 20 minutes at 37°C. Following the EDTA step, small intestines were washed and resuspended in HBSS, vortexed five times for eight seconds each at maximum speed and then washed again in HBSS containing calcium and magnesium (Thermo Fisher, 14025126).

#### Generation of single cell suspensions by enzymatic digestion

Extracted tissues were further processed by several methods depending on the experiment. For all experiments except for *Rag1* fate mapping, tissues were subjected to enzymatic digestion. For *Rag1* fate mapping experiments, tissues were mechanically processed on ice (discussed in the following section). Dura maters, spleens, calvarial bones, leg bones (cleaned of muscle), livers, and small intestines were all chopped in RPMI-H using small scissors. Brain-leptomeninges were placed in dissecting dishes with approximately 200 μL of RPMI-H and minced using razor blades (VWR, 55411-050) until they resembled a fine paste. Each of these tissues was then pelleted at 500 x *g* for five minutes at 4°C in a centrifuge with a swinging-bucket rotor and resuspended in the appropriate enzymatic digestion buffer.

One of two distinct enzymatic digestion protocols was used depending on the experiment type, hereinafter referred to as: Collagenase P or Collagenase P + Dispase II. For the Collagenase P protocol, tissues were digested at room temperature for 60 minutes in a mixture of Collagenase P (0.5 mg/mL, Sigma-Aldrich, 11213865001), DNase-1 (250 U/mL, Worthington, LK003172), and RPMI-H. For the Collagenase P + Dispase II protocol, tissues were digested at 37°C for 30 minutes in a mixture of Collagenase P (0.5 mg/mL), Dispase II (0.8 mg/mL, Thermo Fisher, 17105041), and DNAse-1 (250 U/mL). After digestion, tissues were centrifuged at 500 x *g* for five minutes at 4°C, decanted, and resuspended in 1 mL of FACS buffer. Each tissue was then triturated through a P1000 pipette tip until a single cell suspension was achieved and then filtered through a 70 μm cell strainer (CELLTREAT, 229484).

#### Generation of single cell suspensions by mechanical dissociation

For *Rag1* fate mapping experiments, spleens and lymph nodes were processed mechanically. Spleens were dissociated through 70 μM cell strainers in RPMI-H. Lymph nodes were ground between frosted slides with a small amount of RPMI-H, collected in 2 mL of RPMI-H, and then filtered through 70 μM cell strainers.

#### Organ-specific clean up steps

Brain-leptomeninges, spleen, bone marrow, liver, blood, and small intestine samples were subject to additional clean up steps. Brain-leptomeninges, liver, and small intestine samples were transferred to 15 mL conical tubes with additional FACS buffer added to reach a total volume of 2 mL. 5 mL of 25% (w/v) BSA was added to each sample, and samples were mixed and then centrifuged at 1200 x *g* for 10 minutes at 4°C as a debris removal step. The supernatant and debris layer was removed from each sample by aspiration and the cell pellets transferred to new tubes and washed in FACS buffer. Bone marrow and spleen samples were centrifuged, resuspended in ACK lysis buffer for five minutes on ice, and then quenched with 9 mL of HBSS. For experiments assessing stromal cells in the spleen, single cell suspensions from spleens were then depleted of CD45^+^ cells using the Biolegend MojoSort magnet (Biolegend, 480020) and mouse CD45 nanobeads (Biolegend, 480028). Separation was completed following the manufacturer’s instructions with magnets kept on ice to preserve cell viability. Blood samples were lysed twice for five minutes on ice with quenching and wash steps between. After clean up and lysis steps, all samples were centrifuged and then resuspended in FACS buffer.

#### Antibody staining of single cell suspensions

Single cell suspensions were distributed into 96 well U-bottom plates (Thermo Scientific, 262162) (or in some cases 2 mL microcentrifuge tubes) for antibody staining. For some tissues, a predetermined fraction of the sample was used for staining to reduce the staining volume needed. Samples were centrifuged at 500 x *g* for five minutes at 4°C and then resuspended in viability stain/Fc block solution containing Fixable Viability Dye e780 (1:1000, Thermo Fisher, 65-0865-14), anti-mouse CD16/CD32 (clone 2.4G2, 1:50, BD Biosciences 553141), in some cases CellBlox buffer (1:10, Thermo Fisher, B001T06F01), 2 mM EDTA, and HBSS for 10 minutes on ice. An equal volume of antibody cocktail (antibodies diluted in 20% Brilliant Stain Buffer Plus and FACS buffer) was then added to the cell suspension, mixed, and incubated for an additional 20 minutes on ice. Antibodies used are listed in **Table S1**. Samples were then washed twice with FACS buffer. Cells that would subsequently undergo flow cytometric analysis were fixed for 20 minutes at room temperature in 100 μL of IC Fixation buffer (eBioscience, 00-8222-49), washed twice in FACS buffer, and resuspended in FACS buffer. For CARLIN lineage tracing experiments, live cells were transported to the FACS machine.

#### Flow cytometric acquisition and analysis

Depending on the experiment type, samples were acquired either on a FACSSymphony S6 flow cytometer (FACSDIVA version 9.5), or on a Cytek Aurora 5L (SpectroFlo 3.2.1). CARLIN samples were sorted as described in the following section. Before acquisition, samples were filtered into FACS tubes with 35 μm cell strainer caps (Corning, 352235). In cases where we aimed to quantify the absolute number of cells in a sample, we added 50 μL of counting beads (CountBright Absolute Counting Beads, Thermo Fisher Scientific, C36950) before acquisition. Samples were acquired either in their entirety or to reach a predetermined number of cells or counting beads. FCS files were exported from each machine and analyzed with FlowJo software version 10.10.0.

### Lineage tracing using CARLIN

#### Fluorescence activated cell sorting for CARLIN lineage tracing

For CARLIN experiments, cells were sorted on a FACSAria SORP (special order research product) II flow cytometer (FACSDIVA version 8.0.1). Immediately before acquisition, cells were filtered through a 35 μm cell strainer cap into a FACS tube. Upon acquisition, developing B cells were identified as single, live, dump channel negative (negative for GR-1, CD11b, F4/80, CD11c, CD3, NK1.1, and CD200R3) and then CD19^+^B220^+^CD20^-^IgM^-^IgD^-^. Developing B cells were sorted through a 100 μm nozzle into a 2 mL microcentrifuge tube precoated with 0.5% BSA in HBSS. Sorted samples were centrifuged for five minutes at 500 x *g*, decanted, and resuspended in 1 mL of TRIzol (Thermo Scientific, 15596018). Samples were stored at −20°C or −80°C until further processing.

#### Processing of CARLIN samples for bulk RNA sequencing

RNA from B cell samples was isolated by phenol-chloroform extraction and cDNA was prepared using the Superscript III kit (Thermo Scientific, 18080044). Two nested PCRs and an indexing PCR were performed to generate barcoded libraries following a published protocol and primers^54^. Libraries were quantified by Qubit (Qubit dsDNA HS Assay Kit, Thermo Scientific, Q32854; Invitrogen qubit Assay Tubes, Q32856) and Kapa library quantification (KAPA Library Quantification Kit for Illumina Platforms, Roche, KK4824)), diluted, spiked with 20% PhiX v3 control library (Illumina, FC-110-3001), and paired end sequenced on an Illumina Novaseq 3000 with a 250, 6, 250 read configuration.

#### Analysis of CARLIN lineage tracing data

Binary base call (BCL) files generated from CARLIN library sequencing were converted to fastq format using Illumina’s BCL Convert v4.3.6 software. The converted fastq files were aligned with PEAR (Paired-End reAd mergeR) v0.9.11 and then assembled fastq files were provided to the CARLIN MATLAB pipeline^54^ with the bulk RNA 12-umi CFG primer file. Allele tables—describing the frequency with which each unique CARLIN barcode allele appears in tissues sampled from a single mouse —were assembled using a custom MATLAB script utilizing functions and classes included in the CARLIN pipeline. For each allele table, Allele 0 was excluded as it represents the unedited CARLIN allele. Additionally, barcode alleles were excluded from analysis if their total unique motif identifier (UMI) count across all tissues in a single mouse was less than two. Heatmaps and donut plots were used to present a qualitative visualization of the barcode allele count distribution in each tissue at each development stage. Heatmaps were generated using the pheatmap package^97^ and display the UMI counts for the top 300 alleles found in each tissue from a given mouse. Donut plots were made using ggplot2^98^ and were modified to highlight the top five expanded barcode alleles found in the dura and corresponding alleles in other tissues from the same mouse. Barcode sharing across organs was calculated as the percentage of barcode UMI counts in each organ that share an allele sequence with barcodes in B cells from another organ. The results from this analysis are meant to approximate the proportion of B cells in each organ that likely derive from the same progenitor clone as B cells in other organs.

### Single cell RNA sequencing

#### Preparation of tissues for single-cell RNA sequencing

For single-cell RNA sequencing experiments, we prepared dura, brain-leptomeninges, calvarial bone marrow, and tibial bone marrow tissues in a similar manner to that described above for flow cytometric analysis with a few distinctions. First, mice were injected intracardially with anti-CD45-AF488 (clone 30-F11, 0.2 μg/g of body weight, Biolegend, 103122). Second, to prevent *ex vivo* activation of cells during the enzymatic digestion^99^, we transcardially perfused the mice with a cocktail of transcriptional inhibitors including Actinomycin D (5 μg/mL, Sigma, A1410) and Triptolide (10 μM, Sigma, T3652) in RPMI-H. We also conducted subsequent processing steps before the enzymatic digestion in a buffer containing these transcriptional inhibitors and the translational inhibitor Anisomycin (27.1 μg/mL, Sigma, A9789). Third, we digested the tissues for 30 minutes at 37°C using a mixture of Collagenase P (0.5 mg/mL), Neutral Protease/Dispase (0.8 mg/mL, Worthington, LS02104), DNAse-1 (250 U/mL), Actinomycin D (5 μg/mL), Triptolide (10 μM), Anisomycin (27.1 μg/mL), and DyeCycle Ruby (1:500, V10273). After digestion and trituration, resultant single cell suspensions were blocked with anti-CD16/CD32 (1:50) in HBSS + 2mM EDTA for 10 minutes on ice. Samples were then stained with a primary antibody cocktail for 20 minutes on ice. For dura, bone marrow, and blood samples, this cocktail contained: Brilliant Stain Buffer Plus, CD45-BV605 (clone 30-F11, 1:50, Biolegend, 103155) and TER119-PE (clone TER-119, 1:100, Biolegend, 116208) in FACS buffer. Brain samples were stained with CD45-BV605 and P2RY12-PE (clone S16007D, 1:80, Biolegend, 848004) in Brilliant Stain Buffer Plus and FACS buffer.

#### Fluorescence activated cell sorting for single-cell RNA sequencing

For single cell sequencing experiments, cells were sorted on a FACSAria III flow cytometer (FACSDIVA version 8.0.1). Cells were first filtered through a 35 μm cell strainer cap into a FACS tube and then DAPI added to a final concentration of 0.2 μg/ml. Cells were then sorted through a 100 μm nozzle into a 2 mL microcentrifuge tube precoated with 0.5% BSA (Miltenyi Biotech, 130-091-376) in HBSS. For the dura and bone marrow, single, DAPI^-^ (live), DyeCycle Ruby^+^ (metabolically active), and then CD45^+^CD45 i.v.^-^ (extravascular immune) and CD45^-^TER119^-^(non-immune) cells were sorted to achieve approximately a 1:1 ratio. For the brain, single, DAPI^-^(live), DyeCycle Ruby^+^ (metabolically active), CD45^+^CD45 i.v.^-^, P2RY12^+^CD45^int^ (microglia) and P2RY12^-^CD45^+^ (other immune) cells were sorted to achieve a ratio of approximately 1:4 microglia to non-microglia immune cells.

#### Single cell partitioning, library preparation, and sequencing

Single cell RNA sequencing experiments were performed using 10X Genomics 5’ v1.1 single-cell immune profiling kits. Following cell sorting, the cell suspension was transferred to a PCR tube strip and centrifuged at 300 x *g* for 5 minutes at 4°C. Supernatant was removed to leave a volume equivalent to, or less than, the maximum loading volume (37.8μl). The cell pellet was then resuspended via manual pipetting, which was also used to confirm volume. Any samples with less than 37.8μl had nuclease free water (Thermo Scientific, AM9937) added to reach necessary volume. The tube strip was then lightly pulse vortexed to ensure cells were sufficiently mixed before loading onto Chromium Chip G following 10X Genomics user guide.

After single cell encapsulation and barcoding, single-cell libraries for gene expression were generated following the relevant protocols from 10X Genomics. Libraries were then run on the Agilent 2100 Bioanalyzer system using the Agilent High Sensitivity DNA quantification kit (cat. no. 5067-4626) to assess quality before sequencing.

As described in Marsh et al.^99^, we employed a two-step sequencing process to achieve equivalent sequencing depth across libraries. First, libraries were diluted, pooled, and sequenced at equimolar concentrations on Illumina NextSeq500 using an Illumina NextSeq500/550 High Output v.2.5 150-cycle flow cell with the following parameters: Read 1: 26 bp (16-bp cell barcode, 10-bp UMI); Index 1: 8 bp (Illumina i7 sample index); Read 2: 91 bp (transcript insert).

Resultant data was processed with Cell Ranger 5.0 and used to estimate the number of cells captured per sample. Stock libraries were then diluted and pooled again to account for cell number differences and ensure that the subsequent round of sequencing achieved equal sequencing depth per cell in each sample. Full depth sequencing was performed on NovaSeq 6000 using an S2 100-cycle flow cell by the Broad Institute’s Genomics Platform. The same read length parameters were used for this step as described for sequencing on the NextSeq500. Subsequent analyses were performed solely on data generated on the NovaSeq 6000 platform.

#### Single cell RNA sequencing data processing

Data preprocessing (except for CellBender) was performed on the O2 High Performance Compute Cluster at Harvard Medical School. CellBender processing was performed on virtual instance on Google Cloud Platform.

Raw Illumina bcl files for all libraries were demultiplexed using Cell Ranger v.5.0.0 and bcl2fastq v.2.20.0.422 using the ‘mkfastq’ step with default specifications. Gene expression libraries were then jointly processed using ‘multi’ step which performs gene expression quantification (via ‘count’ step) and then utilizes cell calls from ‘count’ to improve cell calling when performing V(D)J clonotype calling. Gene expression quantification was performed using the mm10 genome (“refdata-gex-mm10-2020-A”; reference annotation corresponds to the filtered version of Ensembl v93).

Ambient RNA was then removed from gene expression count matrices using CellBender v0.2 algorithm^100^. Samples were run using nominal false positive rate of 0.01. Sample specific parameters for ‘expected-cells’ and ‘total-droplets-included’ were adjusted based on Cell Ranger results. The number of training epochs was set to 300 and then dynamically adjusted and re-run on per sample basis depending on results.

To remove low quality cells, we employed two complimentary methods. First, we utilized a data-driven approach that simultaneously evaluates multiple cell quality metrics, as previously described^101^. Second, we clustered the data, manually searched for, and removed cell type multiplets. For the first approach, we constructed a quality network where two cells were connected if they had similar quality based on eight different metrics: (1) expression of genes involved in oxidative phosphorylation, (2) expression of mitochondrial genes, (3) expression of ribosomal protein-coding genes, (4) expression of immediate early genes^99^, (5) percentage of total RNA accounted for by the top 50 most highly expressed genes, (6) percentage of total RNA accounted for by long non-coding RNAs, (7) number of unique genes (log₂-transformed), and (8) number of unique UMIs (log₂-transformed). We then clustered the quality network using the shared nearest neighbor and Leiden clustering algorithms from Seurat v5^102^. Cell clusters were evaluated, and those exhibiting an outlier distribution of quality metrics were excluded.

We then further filtered the dataset for cells that expressed > 500 genes, > 0 % mitochondrial RNA content, and <10% mitochondrial RNA content and clustered the entire dataset using a standard Seurat v5 workflow. We visualized this initial clustering using Uniform Manifold Approximation and Projection (UMAP) and identified and separated immune cell populations (*Ptprc/*CD45^+^) from non-immune cell populations. We then reclustered and subsetted the immune object separately to further distinguish cell types (e.g. B cells, macrophages, etc.) and identify additional cell multiplets and low-quality cells. Cell type (e.g. B cell) and cell subtype (e.g. Pro B cell) designations were determined in subsetted objects and then used for downstream analyses. After performing this process on different subtypes of immune cells, cleaned objects were merged and clustered again to create a final immune cell object. A similar process was performed for the non-immune cell object with a few deviations. We first removed brain-leptomeninges samples from the non-immune dataset, as we did not intend to capture non-immune cells from these samples. We then sequentially subsetted and reclustered cell subtypes from the non-immune object (e.g. FLCs, mesenchymal stem and progenitor cells, etc.), allowing removal of multiplets and finer resolution of cell subtypes. In some cases, we split a given cell type object by tissue (e.g. FLCs isolated from dural samples) for further analysis and subsetting. For FLCs, we also stratified cells into those that likely reside in the dura itself and those that may be contributed by contaminating leptomeninges or other border tissues. This was done by comparing gene expression patterns for FLC clusters in our dataset to those identified in leptomeningeal and brain perivascular FLCs^86^. In general, these putative leptomeningeal FLCs clustered distinctly from the large group of FLCs we designated as dural FLCs. For most analyses of FLCs, we used only the group of FLCs that we designated as dural FLCs.

#### Visualization and analysis of single cell RNA sequencing data

Genes identifying cell clusters were determined using the FindAllMarkers or FindMarkers functions with the default Wilcoxon Rank Sum test in Seurat v5 and selected subsets of markers were plotted as feature plots or dot plots using scCustomize (version 3.1.0)^103^. scCustomize was also used to visualize UMAPs and related plots.

Differentially expressed genes upregulated in dural FLCs were determined using the FindMarkers function in Seurat v5 (parameters: Wilcoxon Rank Sum test, logfc.threshold = 0.585), comparing FLCs to other groups of non-immune cells in the dura or bone marrow compartments. The resulting gene list was filtered for genes passing an adjusted p-value cutoff of 0.05. The percentage of FLCs or cells in the comparator group expressing the given gene was calculated using the Percent_Expressing function from scCustomize v3.0.1 and used to further filter the list for genes expressed in at least 70% of dural FLCs and then generate a specificity score (specificity = % FLCs expressing gene X/ % of cells in comparator group expressing gene X). Genes were then ranked based on their specificity score to prioritize candidates that could be used to target dural FLCs independently of related stromal cell types in the bone marrow. The expression of candidate genes was also evaluated in mouse steady-state scRNA-seq data downloaded from the FibroXplorer database^66^ and plotted as the percentage of a given fibroblast cell type expressing the candidate gene at a level > 0.

#### Ligand-Receptor analysis

Ligand-Receptor analysis was performed to identify interactions between non-immune cells and B cells using the LIANA Rank Aggregate method in the LIANA+ python package^104^ with the “mouseconsensus” resource. This analysis was performed between non-immune cell subtypes (e.g. FLC, OLC, endothelial, etc.) and B cell subsets (e.g. Pro B cell, Mature B cell, etc.), specifically assessing interactions where non-immune cell ligands targeted B cell receptors. Analysis was restricted to cell groups containing 10 or more cells and to ligands or receptors expressed in at least 10% of a given cell group. Ligand-receptor interactions across methods were aggregated using Dimitrov et al.’s implementation of the Robust Rank Aggregation (RRA) originally established by Kolde et al^105^. The UpSetR package^106^ was used to visualize the distribution of ligand-receptor interactions shared across tissues.

## Statistical analyses

With exception of those already described for data derived from sequencing experiments, statistical analyses were conducted in Prism 10. This includes data from flow cytometry and imaging experiments, and comparison of CARLIN barcode sharing across organs. The tests used are specified in figure legends corresponding to each dataset.

## Reagents—antibodies and RNAscope probes

**Table S1.** Antibodies used for flow cytometry and immunohistochemistry. i.v.—used for intravenous labeling. Secondary antibodies (Ab.).

| Target | Fluorophore | Clone | Dilution | Manufacturer | Catalog # |
| --- | --- | --- | --- | --- | --- |
| αSMA | Alexa Fluor 750 | 1A4 | 100 | R&D Systems | IC1420S-025 |
| B220 | Alexa Fluor 647 | RA3-6B2 | 400 | BioLegend | 103229 |
| B220 | Brilliant Violet 480 | RA3 6B2 | 100 | BD Biosciences | 565631 |
| B220 | Brilliant Violet 510 | RA3-6B2 | 80 | BioLegend | 103248 |
| B220 | Brilliant Violet 711 | RA3-6B2 | 160 | BioLegend | 103255 |
| B220 | PE/Cy5.5 | RA3-6B2 | 320 | Thermo Scientific | 35-0452-80 |
| BST-1/CD157 | Brilliant Violet 421 | BP-3 | 80 | BD Biosciences | 740085 |
| BST-2 | Pacific Blue | 129c1 | 400 | BioLegend | 127108 |
| CD115 | Alexa Fluor 488 | AFS98 | 400 | BioLegend | 135512 |
| CD11b | Brilliant Ultraviolet 496 | M1/70 | 100 | BD Biosciences | 749864 |
| CD11b | Brilliant Ultraviolet 496 | M1/70 | 100 | Thermo Scientific | 364-0112-82 |
| CD11c | Alexa Fluor 488 | N418 | 400 | BioLegend | 117311 |
| CD11c | Brilliant Violet 510 | N418 | 160 | BioLegend | 117353 |
| CD11c | PerCP-Fire 806 | N418 | 80 | BioLegend | 117376 |
| CD127 | Brilliant Violet 421 | A7R34 | 80 | BioLegend | 135027 |
| CD13 | APC | QA19A79 | 80 | BioLegend | 164006 |
| CD150 | Brilliant Violet 711 | TC15-12F12.2 | 160 | BioLegend | 115941 |
| CD161/NK1.1 | Brilliant Violet 650 | PK136 | 80 | BioLegend | 108736 |
| CD19 | Alexa Fluor 647 | 6D5 | 400 | BioLegend | 115522 |
| CD19 | Brilliant Violet 510 | 6D5 | 40 | BioLegend | 115546 |
| CD19 | PE/Cy5.5 | eBio1D3(1D3) | 200 | Thermo Fisher Scientific | 35-0193-82 |
| CD19 | RealBlue 705 | 1D3 (RUO) | 40 | BD Biosciences | 570563 |
| CD1d | Brilliant Violet 421 | 1B1 | 160 | BioLegend | 123527 |
| CD20 | Brilliant Ultraviolet 737 | GOT214A | 40 | BD Biosciences | 752756 |
| CD20 | PE/Cy7 | SA275A11 | 160 | BioLegend | 150420 |
| CD200R3 | Alexa Fluor 488 | Ba13 | 400 | Thermo Scientific | 53-2001-82 |
| CD200R3 | RealBlue 744 | Ba13 | 80 | BD Biosciences | 757549 |
| CD23 | Brilliant Ultraviolet 805 | B3B4 | 80 | Thermo Scientific | 368-0232-82 |
| CD24 | Alexa Fluor 488 | M1/69 | 1000 | BioLegend | 101815 |
| CD24 | Brilliant Violet 786 | M1/69 | 80 | Thermo Scientific | 417-0242-82 |
| CD3 | Alexa Fluor 488 | 17A2 | 100 | BioLegend | 100210 |
| CD3 | Alexa Fluor 594 | 17A2 | 100 | BioLegend | 100240 |
| CD3 | Brilliant Violet 510 | 17A2 | 80 | BioLegend | 100234 |
| CD31 | Unconjugated | Polyclonal | 100 | R&D Systems | AF3628 |
| CD31 | Alexa Fluor 647 | MEC13.3 | i.v. | BioLegend | 102516 |
| CD31 | Brilliant Violet 421 | 390 | i.v. | BioLegend | 102424 |
| CD31 | Brilliant Ultraviolet 805 | MEC13.3 | 100 | BD Biosciences | 741939 |
| CD34 | Brilliant Violet 786 | RAM34 | 10 | BD Biosciences | 742971 |
| CD38 | Brilliant Ultraviolet 395 | 90 | 200 | BD Biosciences | 740245 |
| CD39 | Alexa Fluor 647 | DuHa59 | 50 | BioLegend | 143808 |
| CD39 | PE-Cy7 | DuHa59 | 40 | BioLegend | 143806 |
| CD4 | Brilliant Violet 510 | GK1.5 | 80 | BioLegend | 100449 |
| CD4 | Red 718 | RM4-5 | 160 | BD Biosciences | 566939 |
| CD43 | APC | S7 | 160 | BD Biosciences | 560663 |
| CD45 | Alex Fluor 488 | 30-F11 | i.v. | BioLegend | 103122 |
| CD45 | Brilliant Ultraviolet 496 | 30-F11 | 200 | BD Biosciences | 569673 |
| CD45 | Brilliant Ultraviolet 805 | 30-F11 | 200 | BD Biosciences | 568336 |
| CD45 | Brilliant Violet 421 | 30-F11 | 100 | BioLegend | 103134 |
| CD45 | Brilliant Violet 605 | 30-F11 | 100 | BioLegend | 103155 |
| CD45 | Brilliant Violet 785 | 30-F11 | 100 | BioLegend | 103149 |
| CD45 | PE | 30-F11 | i.v. | BioLegend | 103106 |
| CD48 | Brilliant Violet 421 | HM48-1 | 160 | BioLegend | 103428 |
| CD5 | Brilliant Violet 711 | 53-7.3 | 160 | BioLegend | 100639 |
| CD56 | Brilliant Violet 605 | 809220 | 80 | BD Biosciences | 748097 |
| CD64 | Brilliant Violet 711 | X54-5/7.1 | 100 | BioLegend | 139311 |
| CD8 | StarBright Blue 580 | KT15 | 40 | Bio-Rad | MCA609SBB580 |
| CD8a | Brilliant Violet 510 | 53-6.7 | 80 | BioLegend | 100752 |
| CD9 | FITC | MZ3 | 100 | BioLegend | 124808 |
| CD90.2 | Alexa Fluor 488 | 30-H12 | 400 | BioLegend | 105316 |
| CD90.2 | Brilliant Ultraviolet 615 | 53-2.1 | 80 | Thermo Scientific | 366-0902-82 |
| CD90.2 | Brilliant Violet 510 | 30-H12 | 160 | BioLegend | 105335 |
| CD93 | PE/Cy7 | AA4.1 | 160 | BioLegend | 136506 |
| cKit | Brilliant Violet 421 | 2B8 | 100 | BioLegend | 105828 |
| cKit | PE/Cy7 | 2B8 | 160 | BioLegend | 105814 |
| CX3CR1 | Alexa Fluor 488 | SA011F11 | 1000 | BioLegend | 149022 |
| CX3CR1 | Brilliant Violet 480 | Z8-50 | 160 | BD Biosciences | 567824 |
| CX3CR1 | Brilliant Violet 510 | SA011F11 | 1000 | BioLegend | 149025 |
| DLK-1 | Unconjugated | Polyclonal | 200 | R&D Systems | AF8277 |
| Isotype ctr.<br>(for DLK-1) | Unconjugated | Polyclonal | 200 | R&D Systems | AB-108-C |
| ER-TR7 | Alexa Fluor 405 | ER-TR7 | 100 | Novus Biologicals | NB100-64932AF405 |
| ER-TR7 | Alexa Fluor 488 | ER-TR7 | 96 | Novus Biologicals | NB100-64932AF488 |
| EVA1 | Brilliant Violet 711 | G9P3-1 | 40 | BD Biosciences | 752401 |
| F4/80 | Alexa Fluor 488 | BM8 | 100 | BioLegend | 123120 |
| F4/80 | Brilliant Violet 510 | BM8 | 80 | BioLegend | 123135 |
| F4/80 | eFluor 570 | BM8 | 100 | Thermo Fisher Scientific | 41-4801-82 |
| FLT3/CD135 | PE | A2F10 | 40 | BioLegend | 135305 |
| GFP | Unconjugated | Polyclonal | 500 | Aves Labs | GFP-1020 |
| GR-1 | Alexa Fluor 488 | RB6-8C5 | 400 | BioLegend | 108417 |
| GR-1 | Brilliant Ultraviolet 395 | RB6-8C5 | 100 | BD Biosciences | 563849 |
| GR-1 | Brilliant Violet 510 | RB6-8C5 | 200 | BioLegend | 108457 |
| IgD | Alexa Fluor 594 | 11-26c.2a | 100 | BioLegend | 405740 |
| IgD | Alexa Fluor 647 | 11-26c.2a | 100 | BioLegend | 405708 |
| IgD | Brilliant Ultraviolet 395 | 11-26c.2a | 100 | BD Biosciences | 564274 |
| IgM | Alexa Fluor 555 | Polyclonal | 500 | SouthernBiotech | 1020-32 |
| IgM | Brilliant Ultraviolet 395 | R6-60.2 | 100 | BD Biosciences | 564025 |
| IgM | Brilliant Violet 605 | II/41 | 100 | BD Biosciences | 743325 |
| IgM | DyLight 755 | Polyclonal | 500 | Thermo Scientific | SA5-10155 |
| IL-7R | Alexa Fluor 647 | A7R34 | 100 | BioLegend | 135019 |
| KI67 | Alexa Fluor 488 | Sol1a5 | 100 | Thermo Fisher Scientific | 53-5698-82 |
| KI67 | eFluor 570 | Sol1a5 | 100 | Thermo Fisher Scientific | 41-5698-80 |
| Ly-6C | Brilliant Violet 785 | HK1.4 | 400 | BioLegend | 128041 |
| Ly-6G | PE-CF594 | 1A8 | 80 | BD Biosciences | 562700 |
| LYVE-1 | Alexa Fluor 488 | 223322 | 100 | R&D Systems | FAB2125G |
| MCAM/CD146 | Brilliant Ultraviolet 395 | ME-9F1 | 100 | BD Biosciences | 740330 |
| MHC II/I-A/I-E | Spark Violet 538 | M5/114.15.2 | 200 | BioLegend | 107671 |
| MHC-II | Alexa Fluor 488 | M5/114.15.2 | 100 | BioLegend | 107616 |
| NG2 | Unconjugated | EPR23976-145 | 500 | Abcam | ab275024 |
| NK1.1 | Alexa Fluor 488 | PK136 | 400 | BioLegend | 108718 |
| NK1.1 | Brilliant Violet 510 | PK136 | 40 | BioLegend | 108738 |
| P2RY12 | APC | S16007D | 160 | BioLegend | 848006 |
| P2RY12 | PE | S16007D | 160 | BioLegend | 848004 |
| PDGFRa | PE/Cy7 | APA5 | 160 | BioLegend | 135912 |
| PDPN | Alexa Fluor 594 | 8.1.1 | 100 | BioLegend | 127414 |
| PDPN | Brilliant Violet 421 | 8.1.1 | 160 | BioLegend | 127423 |
| S100A9 | Alexa Fluor 647 | 2B10 | 100 | BD Biosciences | 565833 |
| Sca-1 | Alexa Fluor 647 | D7 | 400 | BioLegend | 108118 |
| SCA-1 | Brilliant Violet 510 | D7 | 40 | BioLegend | 108129 |
| Siglec-F | Brilliant Violet 605 | E50-2440 | 160 | BD Biosciences | 740388 |
| ST2 | Alexa Fluor 488 | RMST2-2 | 400 | Thermo Scientific | 53-9335-82 |
| TCR β | Brilliant Ultraviolet 563 | H57-597 | 80 | BD Biosciences | 748406 |
| TCR γδ | Alexa Fluor 488 | GL3 | 200 | BioLegend | 118128 |
| TER-119 | Alexa Fluor 488 | TER-119 | 400 | BioLegend | 116215 |
| TER-119 | Brilliant Violet 711 | TER-119 | 160 | BioLegend | 116267 |
| TER-119 | PE | TER-119 | 200 | BioLegend | 116208 |
| Thy1/CD90.2 | Brilliant Violet 480 | 30-H12 | 160 | BD Biosciences | 746840 |
| *Secondary Ab.* |  |  |  |  |  |
| Chicken IgY | Alexa Fluor 488 | Polyclonal | 2000 | Thermo Scientific | A78948 |
| Goat IgG | Alexa Fluor 647 | Polyclonal | 1000 | Thermo Scientific | A32849 |
| Rabbit IgG | Alexa Fluor 555 | Polyclonal | 1000 | Thermo Scientific | A-31572 |
| Rabbit IgG | Alexa Fluor 647 | Polyclonal | 1000 | Thermo Scientific | A32795 |

**Table S2.**
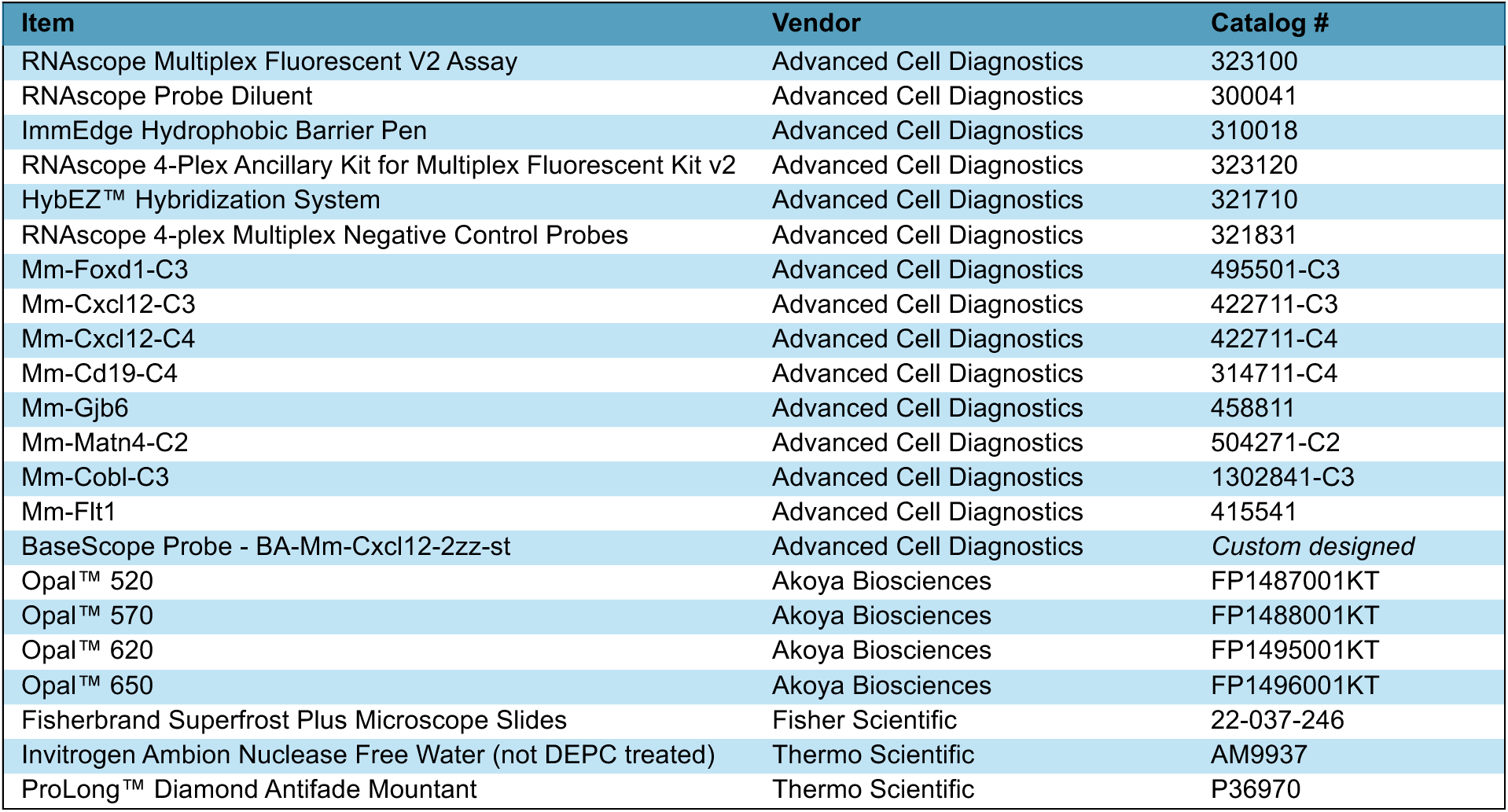
Reagents for RNAscope and BaseScope assays.

| Item | Vendor | Catalog # |
| --- | --- | --- |
| RNAscope Multiplex Fluorescent V2 Assay | Advanced Cell Diagnostics | 323100 |
| RNAscope Probe Diluent | Advanced Cell Diagnostics | 300041 |
| ImmEdge Hydrophobic Barrier Pen | Advanced Cell Diagnostics | 310018 |
| RNAscope 4-Plex Ancillary Kit for Multiplex Fluorescent Kit v2 | Advanced Cell Diagnostics | 323120 |
| HybEZ™ Hybridization System | Advanced Cell Diagnostics | 321710 |
| RNAscope 4-plex Multiplex Negative Control Probes | Advanced Cell Diagnostics | 321831 |
| Mm-Foxd1-C3 | Advanced Cell Diagnostics | 495501-C3 |
| Mm-Cxcl12-C3 | Advanced Cell Diagnostics | 422711-C3 |
| Mm-Cxcl12-C4 | Advanced Cell Diagnostics | 422711-C4 |
| Mm-Cd19-C4 | Advanced Cell Diagnostics | 314711-C4 |
| Mm-Gjb6 | Advanced Cell Diagnostics | 458811 |
| Mm-Matn4-C2 | Advanced Cell Diagnostics | 504271-C2 |
| Mm-Cobl-C3 | Advanced Cell Diagnostics | 1302841-C3 |
| Mm-Flt1 | Advanced Cell Diagnostics | 415541 |
| BaseScope Probe - BA-Mm-Cxcl12-2zz-st | Advanced Cell Diagnostics | *Custom designed* |
| Opal™ 520 | Akoya Biosciences | FP1487001KT |
| Opal™ 570 | Akoya Biosciences | FP1488001KT |
| Opal™ 620 | Akoya Biosciences | FP1495001KT |
| Opal™ 650 | Akoya Biosciences | FP1496001KT |
| Fisherbrand Superfrost Plus Microscope Slides | Fisher Scientific | 22-037-246 |
| Invitrogen Ambion Nuclease Free Water (not DEPC treated) | Thermo Scientific | AM9937 |
| ProLong™ Diamond Antifade Mountant | Thermo Scientific | P36970 |

## RESOURCE AVAILABILITY

### Lead contact

Requests for additional information or resources should be directed to, and will be addressed by, the lead contact Beth Stevens.

### Materials availability

This study did not generate new, unique materials.

### Data and code availability

This preprint will be updated with data deposition links when available.

Any additional information required to analyze the data from this paper will be shared by the lead contact upon request.

## ACKNOWLEDGEMENTS

Joel Cuadrado, Arnaud Frouin, and the Boston Children’s Hospital ARCH staff assisted in managing our mouse colony and performed mouse husbandry. Ron Mathieu at the Boston Children’s Hospital-Harvard Stem Cell Institute flow cytometry core, and Kenneth Ketman, Natasha Barteneva, Ashima Agarwal, and Jodene K. Moore at the Boston Children’s Hospital Flow and Imaging Cytometry Resource Facility provided critical support for flow cytometry and fluorescence activated cell sorting experiments. Chrysothemis Brown at Memorial Sloan Kettering Cancer Center generously provided the *Rag1*^RFP-CreERT2^ mouse strain. Julie Siegenthaler at the University of Colorado Anschutz Medical Campus provided invaluable input about identification of meningeal fibroblast subtypes. Numerous Stevens lab colleagues provided helpful discussion and suggestions throughout the course of this project. Daniel Wilton additionally provided insightful feedback on several versions of this manuscript. We thank each of these individuals for their contributions.

## AUTHOR CONTRIBUTIONS

A.J.W and B.S. conceptualized the study. A.J.W. conducted the majority of experiments and data analysis with support from co-authors. A.J.W., T.M.F, S.K.C, A.S.G, S.E.M., Y.d.S, B.S., A.C.W, Y.H., A.V.M., H.J.B, T.B.L, S.M, and L.D.O contributed to wet lab experiments. A.J.W, J.M.C-R, S.E.M, Y.d.S, V.G. contributed to computational analysis of single cell sequencing and/or lineage tracing data. T.M.F. and J.M.C-R. developed the MATLAB pipeline/GUI for spatial quantification and heatmapping of immune cell localization in whole mount dura mater. A.S.G. developed the modified protocols for RNAscope and BaseScope in whole mount dura mater. F.D.C provided the CARLIN and *Flt3*^EGFP-CreER^ mice, and F.D.C and S.B consulted on implementation of CARLIN lineage tracing experiments. A.J.W. and B.S. wrote the manuscript with input from co-authors, including assistance with figure illustration from L.D.O.

## Supplemental Figures

**Figure S1.**
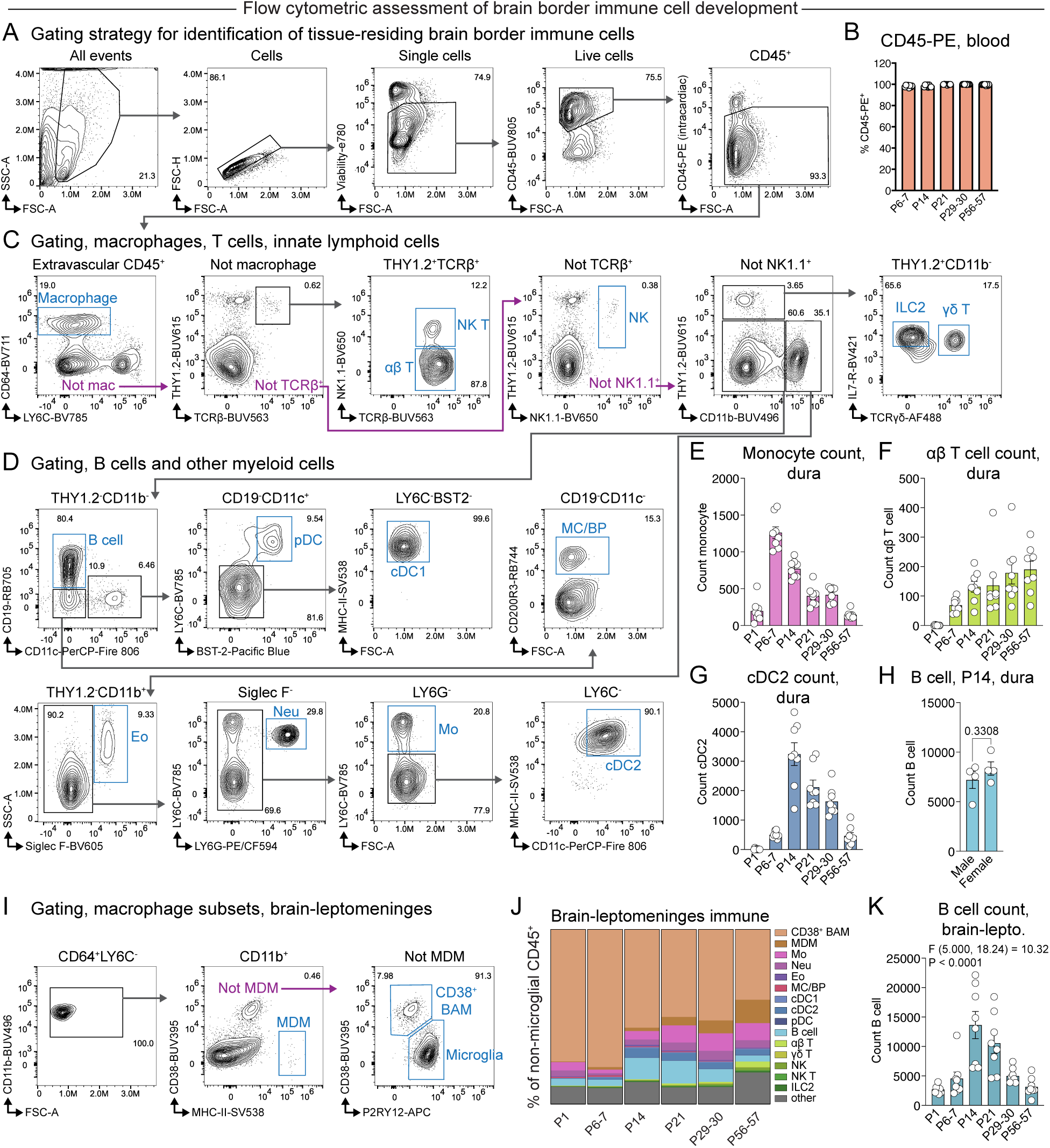
Temporal dynamics of brain border immune compartments in early postnatal life. (A) Flow cytometric gating strategy for assessing the composition of the dural and pooled brain and leptomeningeal immune compartments at different postnatal ages. Live, single cells were identified and then further gated for immune cells expressing CD45. Cells labeled by an intracardiac injection of CD45-PE were then excluded, identifying immune cells present in the extravascular spaces of these tissues for downstream analysis. An example from the P14 dura is shown here; other ages and brain-leptomeningeal samples were gated in a similar manner. (B) Percentage of peripheral blood immune cells labeled by intracardiac injection of CD45-PE. n = 8 C57BL/6 mice per age pooled across two independent experiments for each timepoint. Note that P1 mice were not injected with CD45-PE due to technical limitations and are not shown here. Error bars represent SEM. Welch’s ANOVA: F (4.000, 15.53) = 10.29, P = 0.0003. (C) Gating of macrophages, T cells, and innate lymphoid cells. Extravascular CD45^+^ immune cells were further subsetted based on combinatorial expression of surface markers. Mac—macrophage; αβ T cell; NK T—natural killer T cell; NK—natural killer cell; ILC2—type 2 innate lymphoid cell; γδ T cell. (D) Gating of B cells and other myeloid cells. Cells lacking markers of macrophages, T cells, and innate lymphoid cells were further subsetted based on combinatorial expression of surface markers. THY1.2^-^CD11b^-^ cells (top row): B cells; pDC—plasmacytoid dendritic cells; cDC1—type 1 conventional dendritic cells; MC/BP—mast cells/basophils. THY1.2^-^CD11b^+^ cells (bottom row): Eo—eosinophils; Neu—neutrophils; Mo—monocytes; cDC2—type 2 conventional dendritic cells. (E) Count of dural monocytes across postnatal ages. n = 8 C57BL/6 mice per age pooled across two independent experiments for each timepoint. Error bars represent SEM. Welch’s ANOVA: F (5.000, 18.64) = 62.02, P < 0.0001. (F) Count of dural αβ T cells across postnatal ages. n = 8 C57BL/6 mice per age pooled across two independent experiments for each timepoint. Error bars represent SEM. Welch’s ANOVA: F (5.000, 16.35) = 33.66, P < 0.0001. (G) Count of dural cDC2s across postnatal ages. n = 8 C57BL/6 mice per age pooled across two independent experiments for each timepoint. Error bars represent SEM. Welch’s ANOVA: F (5.000, 16.62) = 68.63, P < 0.0001. (H) Count of dural B cells in male and female mice at P14. n = 4 C57BL/6 mice per sex pooled across 2 independent experiments for each timepoint (from the data presented in Figure 1C). Error bars represent SEM. Student’s T test, unpaired, two-tailed. (I) Subsetting of pooled brain and leptomeningeal macrophage populations. CD64^+^LY6C^-^ macrophages were subdivided into: MDM—monocyte–derived macrophages; CD38^+^ BAMs—border-associated macrophages; and microglia. (J) Quantification of flow cytometry data depicting immune cell proportions in the pooled brain and leptomeninges at postnatal developmental timepoints. Microglia are excluded from this analysis to allow better visualization of changes in other immune subtypes. Each value represents the mean of n = 8 C57BL/6 mice per timepoint collected across two independent experiments for each timepoint. CD38^+^ BAM—border-associated macrophage; MDM— monocyte-derived macrophage; Mo—monocyte; Neu—neutrophil; Eo—eosinophil; MC/BP—mast cell/basophil; cDC1—type 1 conventional dendritic cell; cDC2—type 2 conventional dendritic cell; pDC—plasmacytoid dendritic cell; B cell; αβ T cell; γδ T cell; NK—natural killer cell; NK T—natural killer T cell; ILC2—type 2 innate lymphoid cell; other—uncategorized CD45^+^ cells. (K) Quantification of flow cytometry data depicting total CD19^+^ B cell counts in pooled brain and leptomeninges (brain-lepto.) at postnatal developmental timepoints. n = 8 mice C57BL/6 mice per timepoint collected across 2 independent experiments for each timepoint. Error bars represent SEM. Welch’s ANOVA.

**Figure S2.**
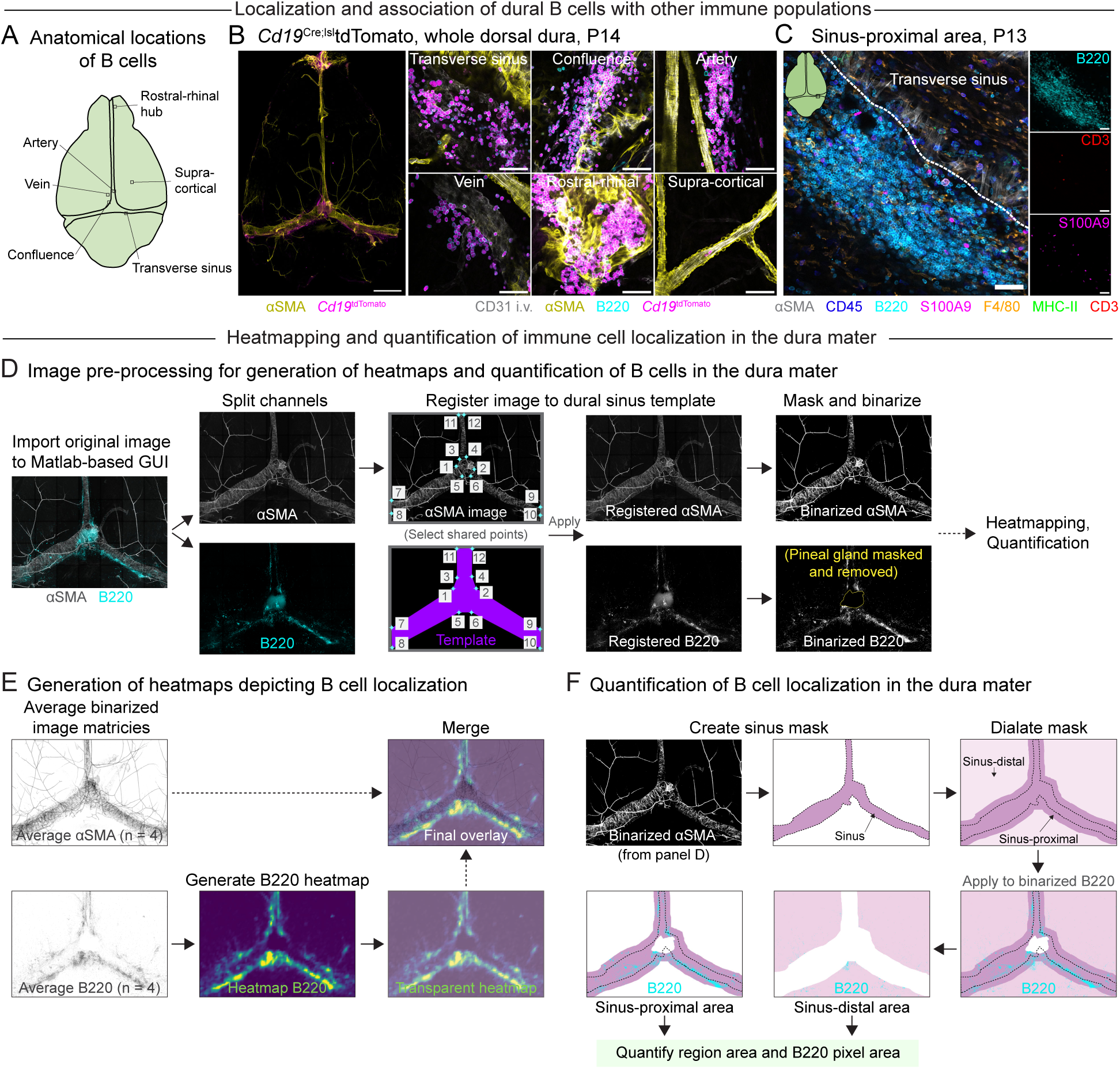
Spatial mapping of dural immune cells in the early-life window. (A) Diagram of a whole mount preparation of the dorsal cranial dura. Areas of interest corresponding to images in (B) are highlighted. (B) Representative tile scan image of the dorsal dura in a P14 *Cd19*^Cre^;^lsl^tdTomato mouse (left). High magnification images (right) depict B cell clusters in various regions of the dura. CD31 i.v.—blood vessels, white; `α`SMA—α-smooth muscle actin, yellow; B220—B cells, cyan; *Cd19*^tdTomato^—B cells, magenta. Scale bar for tile scan image indicates 500 µm. Scale bars for high magnification images indicate 50 µm. (C) Representative high magnification image of a sinus-proximal B cell cluster in the C57BL/6 mouse dura at P13. `α`SMA—α-smooth muscle actin, white; CD45—immune cells, blue; B220—B cells, cyan; S100A9—neutrophils, magenta; F4/80—macrophages, orange; MHC-II—antigen presenting cells, green; CD3—T cells, red. The edge of the sinus is marked with a white dotted line. Scale bars indicate 50 µm. (D) Image pre-processing steps implemented in a custom MATLAB GUI that allows heatmapping and quantification of dural immune cells from tile scan images. Z-projected tile scan images including `α`SMA (structural marker of the dural sinuses) and B220 (B cell marker) were first split into individual channels. The `α`SMA channel was registered to a template of the dural sinuses and these registration coordinates were applied to the B220 channel, allowing multiple images to be overlaid in the subsequent heatmapping step. Images were then binarized and autofluorescent signal from the pineal gland masked and removed before downstream processing. (E) To generate heatmaps of B cell localization, registered, binarized images from multiple mice were merged and averaged per channel. The averaged B220 image was used to make a heatmap describing the average occurrence of B cell signal at each X-Y coordinate and overlaid with the averaged `α`SMA image to generate the final heatmap. The final heatmap shown here is derived from Figure 1G and is included again to allow illustration of the entire image processing pipeline. (F) Registered and binarized images were also used to quantify the amount of B cell signal in different regions of the dura. The binarized `α`SMA channel (image derived from panel D) was used to generate masks of the sinus-proximal area (overlying and within 300 µm of the outer edge of the sinuses) and the sinus-distal area (> 300 µm from the edges of the sinuses). These masks were then applied to the B220 channel and used to calculate the total area of each region and the area of each region occupied by B220 signal.

**Figure S3.**
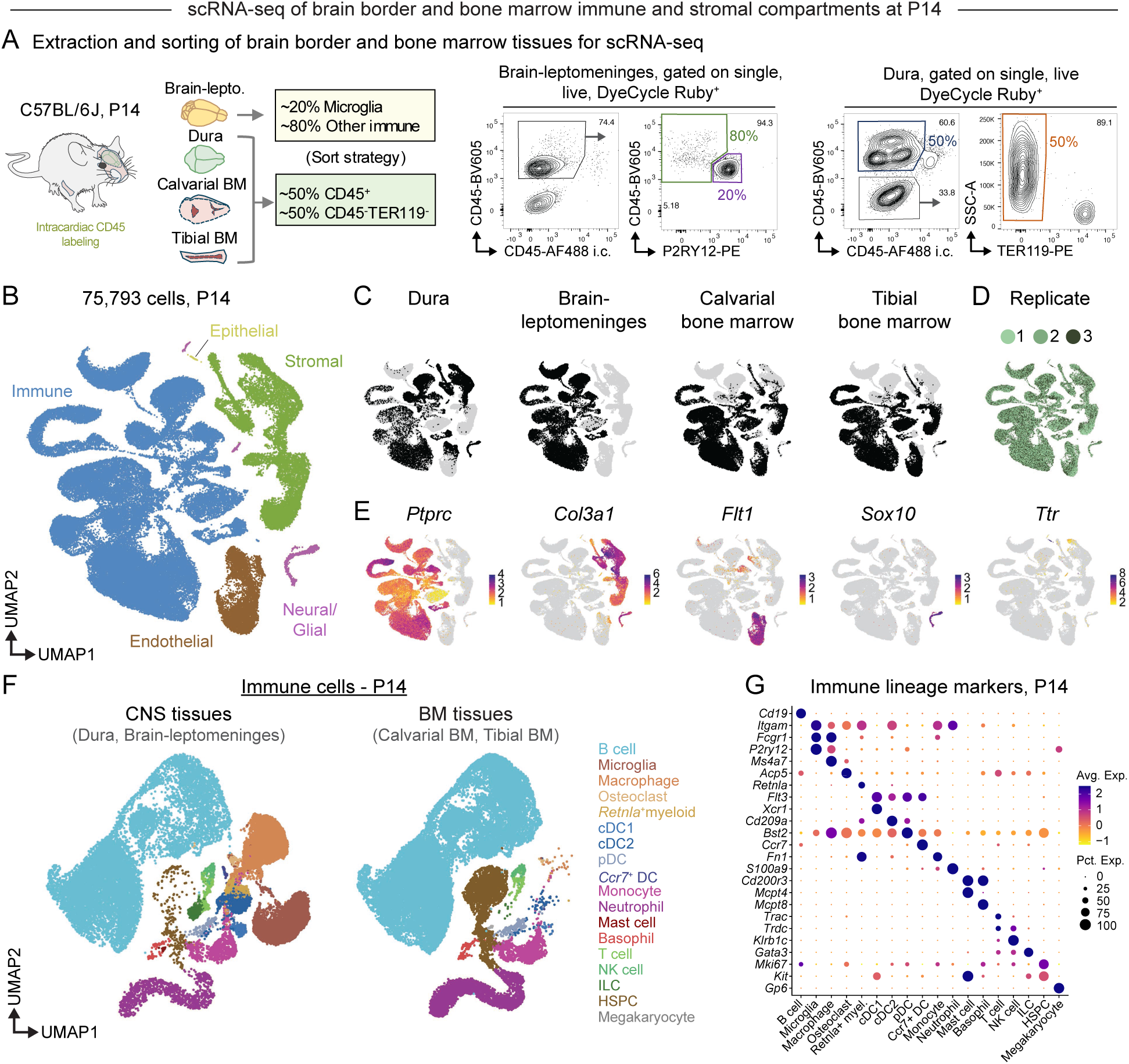
Molecular profiling of early-life brain border and bone marrow compartments. (A) Experimental schematic and gating strategy—fluorescence activated cell sorting of immune and non-immune cells for scRNA-seq. P14 mice were labeled intracardially (i.c) with CD45-AF488 and transcardially perfused. Dura, pooled brain and leptomeninges (Brain-lepto.), calvarial bone marrow (BM), and tibial bone marrow were extracted, digested into single cell suspensions, stained, and sorted to capture extravascular CD45^+^ cells from all organs and CD45^-^TER119^-^ cells from the dura, calvarial bone marrow, and tibial bone marrow. In pooled brain and leptomeningeal samples, P2RY12^+^CD45^int^ microglia were down sampled to allow better resolution of non-microglial cell types. Gating strategy for brain-leptomeninges: single cells, DAPI^-^ (live), DyeCycle Ruby^+^ (metabolically active), CD45^+^CD45 i.c.^-^, P2RY12^+^CD45^int^ (microglia—sorted) and P2RY12^-^CD45^+^ (other immune— sorted). Gating strategy for dura, calvarial bone marrow, and tibial bone marrow: single cells, DAPI^-^ (live), DyeCycle Ruby^+^ (metabolically active), and then CD45^+^CD45 i.c.^-^ (immune—sorted) and CD45^-^TER119^-^ (non-immune— sorted). Cells were then captured on the Chromium 10x platform for scRNA-seq. (B) UMAP representation of all cells across organs sampled at P14 from three male mice. Major cell classes are highlighted. (C) Distribution of cells across organs. (D) Distribution of cells across mouse replicates. (E) Markers defining cell classes. *Ptprc* (CD45)—immune cells; *Col3a1*—stromal cells; *Flt1*—endothelial cells; *Sox10*—Schwann cells/myelinating cells, part of neural/glial grouping; *Ttr*—choroid plexus epithelial cells. (F) UMAP representation of immune cells, split by CNS (dura and brain-leptomeninges) and BM (calvarial and tibial bone marrow) tissues. Cells are grouped into major immune cell types. cDC1—type 1 conventional dendritic cell; cDC2—type 2 conventional dendritic cell; pDC—plasmacytoid dendritic cell; *Ccr7*^+^ DC—*Ccr7*^+^ dendritic cell; NK cell—natural killer cell; ILC—innate lymphoid cell; HSPC—hematopoietic stem and progenitor cell. (G) Marker genes for immune cell types denoted in (F).

**Figure S4.**
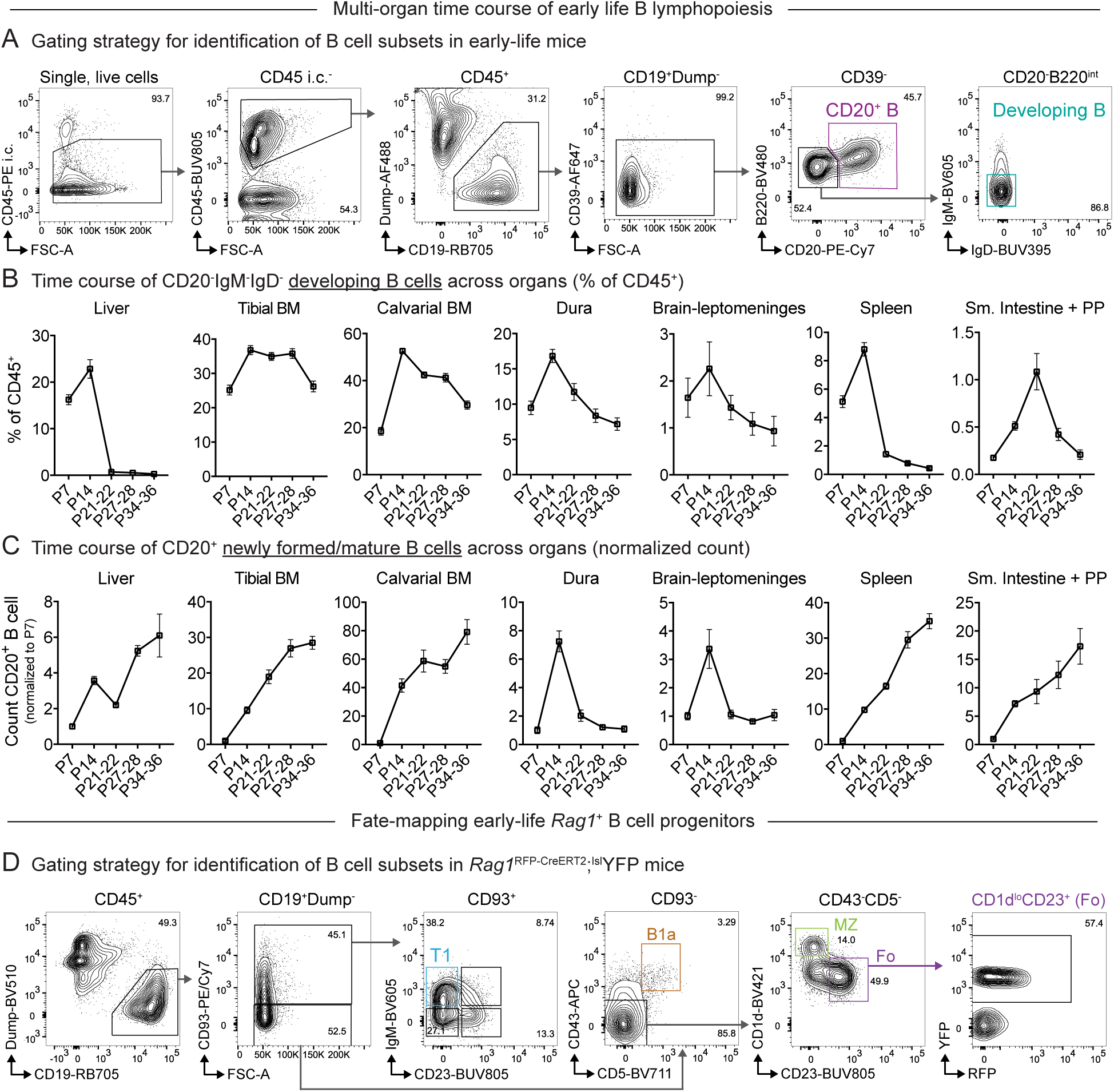
Mapping early life developing B cells and their contribution to the mature B cell compartment. (A) Gating strategy for identifying developing and newly formed/mature B cells over a time course of postnatal development. The P14 dura is shown and is representative of other organs and timepoints examined. B cells were gated as single, live, intracardiac (i.c.) CD45^-^, CD45^+^, CD19^+^/dump channel negative (negative for GR-1, CD115, F4/80, CD11c, CD90.2, CD3, NK1.1, CD200R3, CX3CR1, TER119) cells. Plasma cells were excluded by gating for CD39^lo^ B cells, and then developing B cells identified as B220^int^CD20^-^, IgM^-^IgD^-^. Newly formed/mature B cells were identified as CD20^+^ cells from the CD39^lo^ gate. Note that the intracardiac CD45 gate was not used for liver samples due to substantial leakage of the CD45 antibody into the tissue. (B) Developing B cells as a percentage of CD45^+^ cells in primary lymphoid and extramedullary organs across a time course of postnatal development, assessed by flow cytometry. n = 8 C57BL/6 mice per age at P7, P27-28, and P34-36, and 12 mice per age at P14 and P21-22, pooled from 2-3 independent experiments per age. Error bars indicate SEM. Welch’s ANOVA. Liver: F(4.000, 19.96) = 93.36, P < 0.0001; Tibial bone marrow (BM): F(4.000, 20.04) = 14.21, P < 0.0001; Calvarial bone marrow: F(4.000, 19.21) = 85.26, P < 0.0001; Dura: F(4.000, 20.67) = 15.66, P < 0.0001; Brain-leptomeninges: F(4.000, 20.35) = 1.337, P = 0.2902; Spleen: F(4.000, 20.36) = 99.28, P < 0.0001; Small intestine + Peyer’s patches (PP): F(4.000, 20.14) = 15.25, P < 0.0001. (C) Normalized count of CD20^+^ newly formed/mature B cells from primary lymphoid and extramedullary organs across a time course of postnatal development, assessed by flow cytometry. Numbers of B cells for each organ were normalized to the mean B cell count at P7. n = 8 C57BL/6 mice per age at P7, P27-28, and P34-36, and 12 mice per age at P14 and P21-22, pooled from 2-3 independent experiments per age. Welch’s ANOVA. Liver: F(4.000, 19.35) = 217.2, P < 0.0001; Tibial bone marrow (BM): F(4.000, 20.17) = 59.95, P < 0.0001; Calvarial bone marrow: F(4.000, 20.42) = 48.23, P < 0.0001; Dura: F(4.000, 20.81) = 12.81, P < 0.0001; Brain-leptomeninges: F(4.000, 20.14) = 7.723, P = 0.0006; Spleen: F(4.000, 20.12) = 138.9, P < 0.0001; Small intestine + Peyer’s patches (PP): F(4.000, 19.36) = 6.115, P = 0.0024. (D) Gating strategy for identifying transitional and mature B cell subsets in *Rag1*^RFP-CreERT2^;^lsl^YFP fate mapping experiments. A representative example from the spleen is shown for the P24 harvest timepoint. B cells were identified as single, live, CD45^+^, CD19^+^/dump channel negative (negative for GR-1, F4/80, CX3CR1, CD11c, CD3, CD4, CD8a, NK1.1). CD93^+^ and CD93^-^ B cells were then separated. Transitional T1 B cells were identified from the CD93^+^ fraction as IgM^+^CD23^-^. From the CD93^-^ gate, B1a B cells were identified as CD43^+^CD5^+^ and B2 B cells were gated as CD43^-^CD5^-^. Marginal zone (MZ) B cells were identified as CD1d^hi^CD23^-^ and follicular B cells as CD1d^lo^CD23^+^. A representative example of YFP and RFP expression in splenic follicular B cells is shown. Inguinal lymph nodes were gated by the same strategy except that only follicular B cells were identified and assessed for YFP expression.

**Figure S5.**
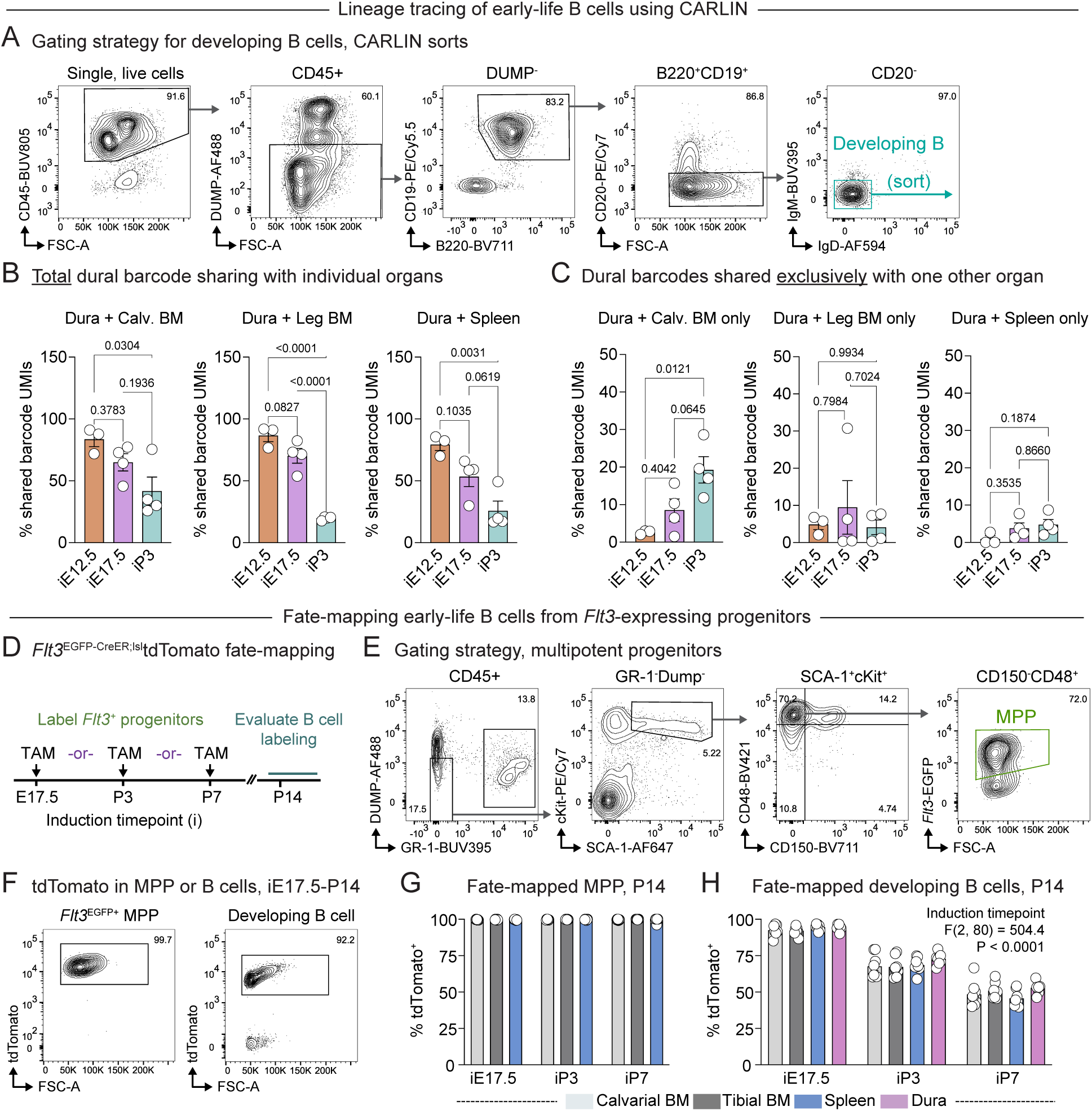
Lineage tracing and fate-mapping of early-life B cell progenitors. (A) Gating strategy used to sort developing B cells for CARLIN experiments. B cells were gated as live, single, CD45^+^, Dump channel negative (negative for GR-1, F4/80, CD11c, CD3, NK1.1, CX3CR1, CD200R3, ST2, TER119), B220^+^CD19^+^ cells. Developing B cells (representing pooled Pro and Pre B cell stages) were further gated as CD20^-^, IgM^-^IgD^-^ cells and sorted for downstream RNA sequencing. Gating strategy shown is from leg bone marrow and is representative of other organs. (B) Total percentage of dural B cell CARLIN barcode UMIs that share an allele sequence with B cell barcodes in calvarial bone marrow (Calv. BM), leg bone marrow, or spleen. n = 3 mice at iE12.5, 4 mice at iE17.5, and 4 mice at iP3. Error bars represent SEM. One-way ANOVA with Tukey’s multiple comparisons test. Dura-Calvarial BM: F(2, 8) = 5.239, P = 0.0351; Dura-Leg BM: F(2, 8) = 61.12, P < 0.0001; Dura-Spleen: F(2, 8) = 12.01, P = 0.0039. (C) Percentage of dural B cell CARLIN barcode UMIs that share an allele sequence exclusively with B cell barcodes in one other organ. n = 3 mice at iE12.5, 4 mice at iE17.5, and 4 mice at iP3. Error bars represent SEM. One-way ANOVA with Tukey’s multiple comparisons test. Dura-Calvarial BM: F(2, 8) = 7.913, P = 0.0127; Dura-Leg BM: F(2, 8) = 0.3816, P = 0.6945; Dura-Spleen: F(2, 8) = 1.992, P = 0.1985. (D) Experimental design for evaluating FLT3^+^ progenitor contribution to early-life B cells. *Flt3*^EGFP-CreER^;^lsl^tdTomato mice were induced with tamoxifen at iE17.5, iP3, or iP7, and then B cells evaluated at P14 for tdTomato labeling. (E) Gating strategy for identification of multipotent progenitors (MPP) used as a reference for labeling efficiency in *Flt3*^EGFP-CreER^;^lsl^tdTomato fate mapping experiments. MPPs were identified as live, single, CD45^+^, Dump channel negative (negative for CD19, B220, F4/80, CD11c, CD3, CD4, CD8a, NK1.1)/GR-1^-^, SCA-1^+^cKit^+^, CD150^-^CD48^+^, *Flt3*^EGFP+^. Gating shown for calvarial bone marrow and is representative of other organs. (F) Representative flow cytometry contour plots showing tdTomato expression in multipotent progenitors (MPP) or developing B cells in calvarial bone marrow of a *Flt3*^EGFP-CreER^;^lsl^tdTomato mouse induced at iE17.5 and evaluated at P14. MPPs were gated as shown in (E). Developing B cells were gated as CD45^+^, CD19^+^Dump channel-negative (negative for GR-1, CD11c, CD3, CD4, CD8a, NK1.1), CD20^-^, IgM^-^IgD^-^, representing Pro and Pre B cell populations. (G) Percentage of tdTomato^+^ MPPs from calvarial bone marrow, tibial bone marrow, or spleen in *Flt3*^EGFP-CreER^;^lsl^tdTomato mice induced at embryonic or postnatal timepoints and evaluated at P14. n = 7 mice from iE17.5, 8 mice from iP3, and 8 mice from iP7 pooled from two independent experiments for each induction timepoint. Error bars represent SEM. Two-way ANOVA. Induction timepoint: F(2, 60) = 0.6248, P = 0.5388; Tissue: F(2, 60) = 1.723, P = 0.1873; Interaction: F(4, 60) = 1.135, P = 0.3488. (H) Percentage of tdTomato^+^ developing B cells from calvarial bone marrow, tibial bone marrow, spleen, or dura in *Flt3*^EGFP-CreER^;^LSL^tdTomato mice induced at embryonic or postnatal timepoints and evaluated at P14. n = 7 mice from iE17.5, 8 mice from iP3, and 8 mice from iP7 pooled from two independent experiments for each induction timepoint. Error bars represent SEM. Two-way ANOVA. Induction timepoint: F(2, 80) = 504.4, P < 0.0001; Tissue: F(3, 80) = 2.642, P = 0.0549; Interaction: F(6, 80) = 1.072, P = 0.3861.

**Figure S6.**
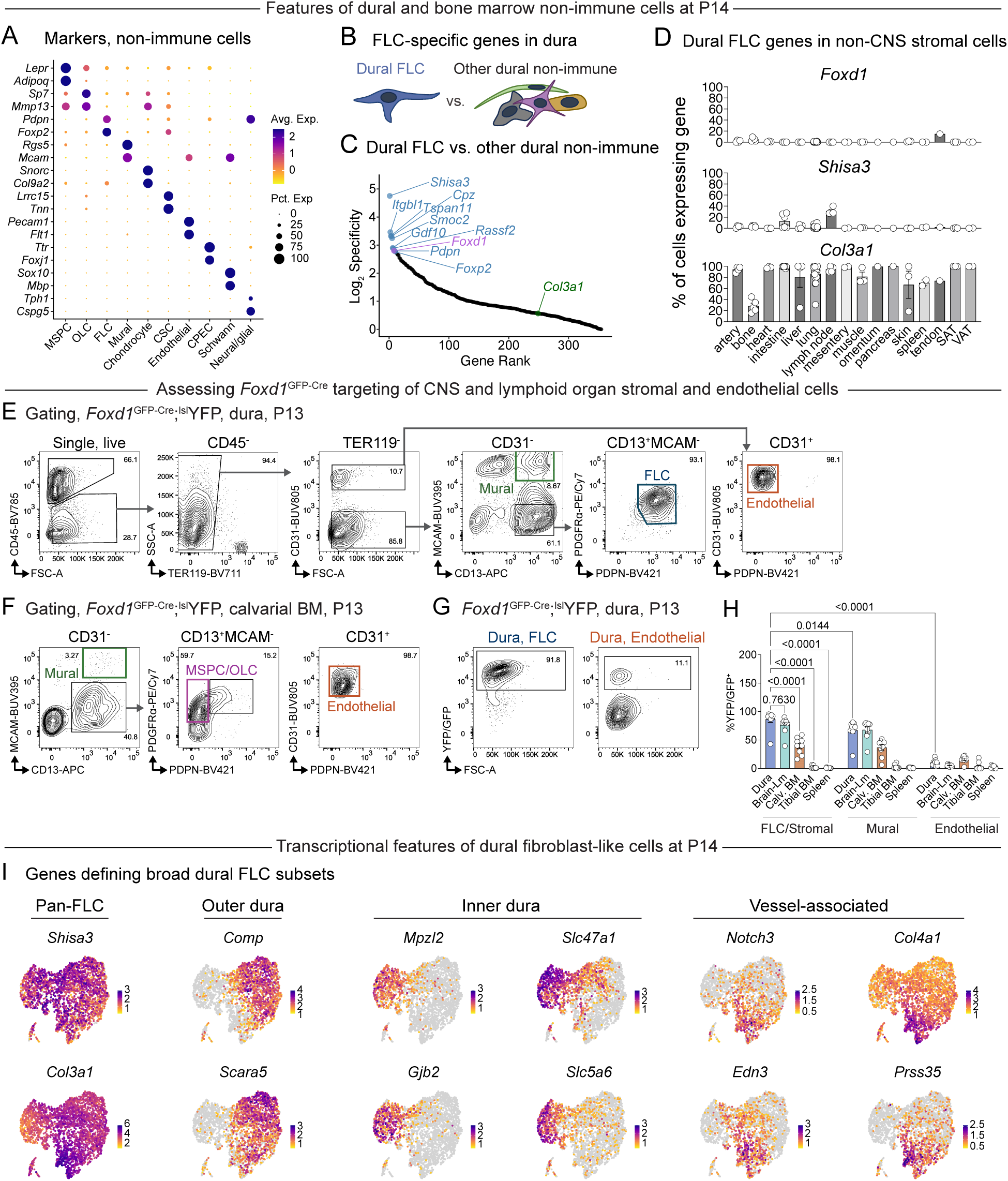
Molecular characterization and targeting of CNS stromal niches. (A) Markers defining non-immune cell subtypes across the dura, calvarial bone marrow, and tibial bone marrow at P14 (related to Figure 5A). MSPC—Mesenchymal stem and progenitor cell; OLC—Osteolineage cell; FLC—Fibroblast-like cell; Mural cell (smooth muscle cell and pericyte); Chondrocyte; CSC—Cranial stromal cell; Endothelial cell; Schwann cell; Neural/glial cell (pinealocyte, astrocyte-like cell, neuronal cell); CPEC—Choroid plexus epithelial cell. (B) Experimental design for identification of genes enriched in and specific for dural FLCs compared to other non-immune cells in the dura. Differentially expressed genes upregulated in dural FLCs were assigned a specificity score where specificity = % expression in FLCs/% expression in other dural non-immune cells. This list was further filtered for genes expressed in at least 70% of dural FLCs. (C) Dural FLC-enriched genes, ranked by specificity and plotted on a log_2_ scale. The top 10 genes specific to dural FLCs are labeled (top 10, blue; gene of interest *Foxd1*, purple), as well as the global fibroblast marker *Col3a1* (green). (D) Expression of dural FLC-enriched genes *Foxd1* and *Shisa3* and global fibroblast marker *Col3a1* in fibroblasts and other stromal populations across the body. Data is derived from FibroXplorer and includes a compilation of scRNA-seq datasets describing fibroblasts across steady state mouse organs. For each organ, the percentage of fibroblasts/stromal cells expressing the candidate gene at a level > 0 was calculated and plotted. SAT— subcutaneous adipose tissue; VAT—visceral adipose tissue. (E) Representative gating strategy to identify stromal and endothelial cell populations in *Foxd1*^GFP-Cre^;^lsl^YFP mice at P13-P14. The dura is shown here and is representative of gating for pooled brain and leptomeninges, and spleen, samples. After gating live, single, CD45^-^, TER119^-^, cells, endothelial cells were identified as CD31^+^PDPN^-^. CD31^-^ cells were divided into CD13^+^MCAM^+^ (mural) and CD13^+^MCAM^-^ groups. In the dura, FLCs were identified from CD13^+^MCAM^-^ group as PDGFR`α`^+^PDPN^+^ cells. In the brain-leptomeninges and spleen, FLCs/stromal cells were designated as CD13^+^MCAM^-^ due to variable expression of PdGfRα and PDPN. (F) Representative gating strategy to identify stromal and endothelial cell populations in *Foxd1*^GFP-Cre^;^lsl^YFP mice at P13-P14. The calvarial bone marrow (BM) is shown here and is representative of tibial bone marrow gating. After gating live, single, CD45^-^, TER119^-^, cells, endothelial cells were identified as CD31^+^PDPN^-^. CD31^-^ cells were divided into CD13^+^MCAM^+^ (mural) and CD13^+^MCAM^-^ groups. CD13^+^MCAM^-^ cells were further gated for PDGFR`α`^+^PDPN^-^ cells to identify MSPC/OLC populations. (G) Representative flow cytometry plots of YFP/GFP expression in dural FLCs and dural endothelial cells P13 in *Foxd1*^GFP-Cre^;^lsl^YFP mice. Note that this fluorescent signal reflects a combination of the recombined YFP allele and some *Foxd1*-driven GFP from the Cre line. (H) Percentage of stromal and endothelial populations from P13-P14 *Foxd1*^GFP-Cre^;^lsl^YFP mice expressing YFP/GFP, indicating targeting by *Foxd1*-driven Cre. Note that this fluorescent signal reflects a combination of the recombined YFP allele and some *Foxd1*-driven GFP from the Cre line. FLC/Stromal (dural FLCs, pooled brain-leptomeningeal (Lm) FLCs, calvarial bone marrow (Calv. BM) and tibial bone marrow MSPCs/OLCs, and spleen stromal cells); MCAM^+^CD13^+^ mural cells from all organs; and CD31^+^PDPN^-^ blood endothelial cells from all organs. n = 9 mice pooled across three independent experiments. Error bars indicate SEM. Two-way ANOVA with Tukey’s multiple comparisons test. Cell type x Tissue interaction: F(8, 120) = 33.10, P < 0.0001. Select post-hoc tests are displayed comparing dural FLCs to other relevant cell types. (I) Additional genes defining broad groups of dural FLCs (outer dura, inner dura, vessel-associated dura), corresponding to Figure 5H.

**Figure S7.**
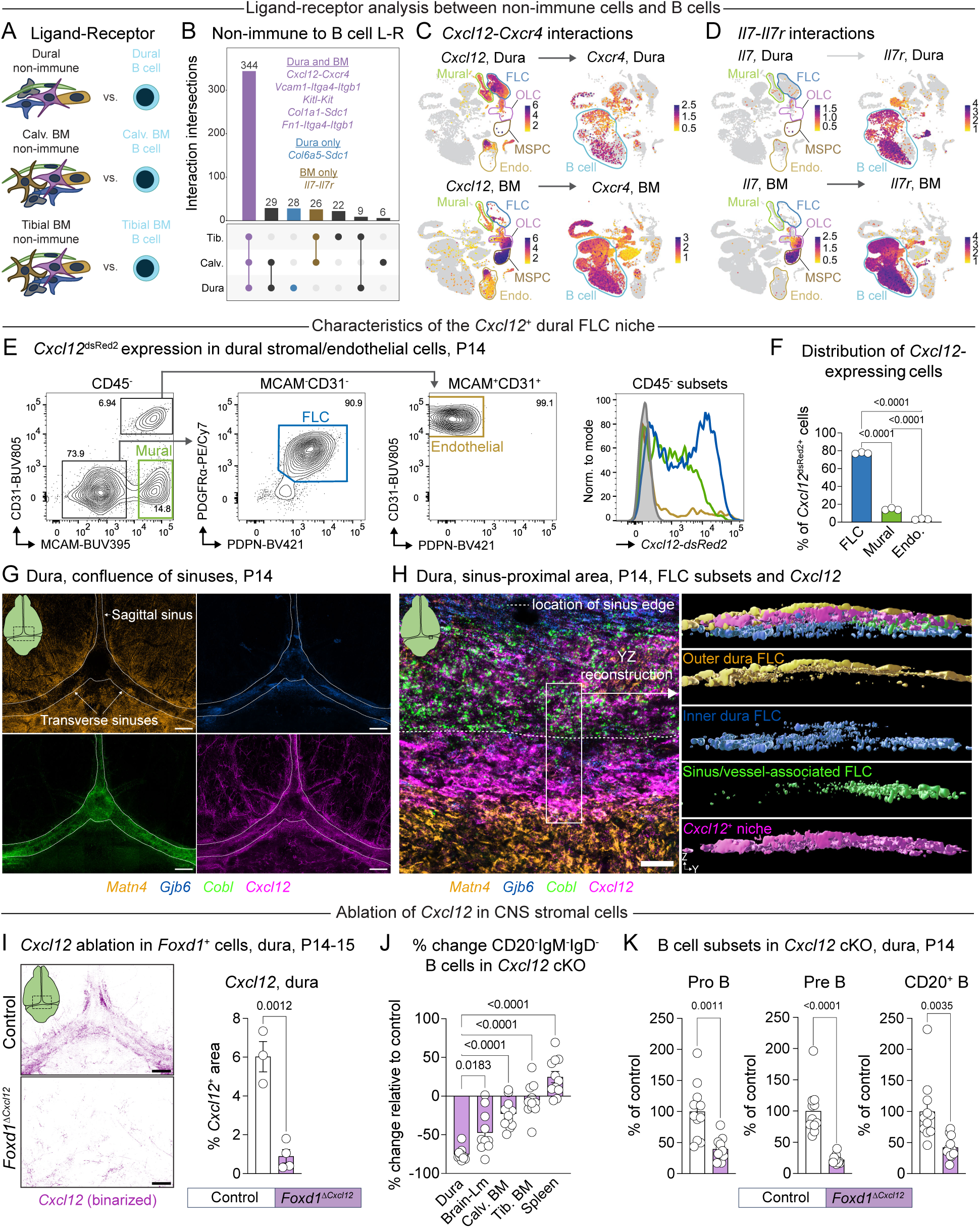
Characterization of lymphopoietic niches in the early-life dura and bone marrow. (A) Ligand-receptor interactions between non-immune cells and B cells were calculated for the dura, calvarial bone marrow (Calv. BM), and tibial bone marrow separately using LIANA. (B) Intersection of ligand-receptor interactions between non-immune cells and B cells in the dura, calvarial bone marrow (Calv.), and tibial bone marrow (Tib.), from analysis described in (A). Numbers of shared versus unique non-immune cell–to–B cell interactions plotted across organs and select shared or unique interactions are highlighted. (C) Expression of *Cxcl12* and cognate receptor *Cxcr4* in dural or bone marrow (BM) non-immune and immune cells at P14. FLC—Fibroblast-like cell; OLC—Osteolineage cell; MSPC—Mesenchymal stem and progenitor cell; Endo.— Endothelial cell; Mural cell (pericyte and smooth muscle cell); B cell. (D) Expression of *Il7* and cognate receptor *Il7r* in dural or bone marrow (BM) non-immune and immune cells at P14. FLC—Fibroblast-like cell; OLC—Osteolineage cell; MSPC—Mesenchymal stem and progenitor cell; Endo.— Endothelial cell; Mural cell (pericyte and smooth muscle cell); B cell. (E) Flow cytometric identification of stromal and endothelial subsets in *Cxcl12*^dsRed2^ mice at P14. Live, single, CD45^-^ cells were further gated to identify endothelial cells (MCAM^+^CD31^+^PDPN^-^), mural cells (MCAM^+^CD31^-^), and FLCs (MCAM^-^CD31^-^PDGFR`α`^+^PDPN^+^). A histogram of *Cxcll2* ^dsRed2^ expression in each cell type is shown compared to CD45^-^ cells from a dsRed2-negative littermate control. DsRed2-negative cells, grey; endothelial cells, gold; mural cells, green; FLCs, blue. (F) Percent of total CD45^-^, *Cxcl12*^dsRed2^ expressing cells in the P14 dura that are FLCs, mural cells, or endothelial cells. n = 3 littermates from one experiment. Error bars indicate SEM. One-way ANOVA with Tukey’s multiple comparisons test. F (2, 6) = 5667, P < 0.0001. (G) Representative confocal tile scan of the confluence of the sinuses at P14 depicting the localization of *Cxcl12* and FLC subsets. *Matn4*—outer dura, orange; *Gjb6*—inner dura, blue; *Cobl*—sinus/perivascular dura, green, *Cxcl12*, magenta. White outlines indicate the approximate location of the dural sinuses based on *Cobl* expression. Image represents a maximum intensity Z projection across the depth of the tissue. Scale bar indicates 500 µm. (H) Localization of *Cxcl12* and dural FLC subsets composing the outer dura (*Matn4*), inner dura (*Gjb6*), and sinus/vessel-associated dura (*Cobl*). Image represents a maximum intensity projection of 3 µm from a confocal Z stack localized to an area near the transverse sinus (left). Scale bar represents 50 µm. The full Z stack was used to reconstruct this region in 3D using Imaris. Surfaces were constructed out of each channel and then used to visualize the location of *Cxcl12* signal relative to FLC subtypes in the YZ dimension (right). (I) Representative tile scan images depicting *Cxcl12* expression at the confluence of the dural sinuses in control (*Cxcl12^ffl/f^*) or *Foxdl*^Δ^*^Cxcl^*^12^ (*Foxdl*^GFP-Cre/+^;*Cxcll2*™) mice at P14 (left). Quantification of *Cxcl12* signal (right). Expression of *Cxcl12* exon 2, which is flanked by loxP sites in this strain, was visualized by BaseScope. *Cxcl12* signal was binarized before quantification. n = 3 Control and n = 4 *Foxdl^ΔCxcf12^* mice from two litters each containing Control and *Foxdl^ΔCxcf12^* pups at P14-15. Error bars indicate SEM. Student’s T test, unpaired, two-tailed. (J) Flow cytometric quantification of developing (CD19^+^CD20^-^IgM^-^IgD^-^) B cells across organs at P14 in *Foxdl^ΔCxcf12^* mice, related to Figure 6F. n = 11 control and 11 *Foxdl^ΔCxcf12^* mice pooled from two independent experiments each consisting of one litter containing Control and *Foxdl^ΔCxcf12^* littermates. Percent change in B cell counts in *Foxdl^ΔCxcf12^* mice relative the Control group is plotted for each organ. Error bars indicate SEM. Welch’s ANOVA with Dunnett’s T3 multiple comparisons test. F (4.000, 23.07) = 67.33, P < 0.0001. Select post-hoc comparisons shown. Brain-Lm—pooled brain and leptomeninges; Calv. BM—calvarial bone marrow; Tib. BM—Tibial bone marrow. (K) Flow cytometric quantification of dural B cell subsets at P14 in control and *Foxdl^ΔCxcf12^* mice. Within CD19^+^CD20^-^ IgM^-^IgD^-^ cells, Pro B cells were identified as cKit^+^CD24^lo^ and Pre B cells as cKit^-^CD24^hi^. CD20^+^ B cells were identified as CD19^+^CD20^+^. n = 11 control and 11 *Foxdl^ΔCxcf12^* mice pooled from two independent experiments each consisting of one litter containing Control and *Foxdl^ΔCxcf12^* littermates. For each experiment, B cell counts were normalized to the mean count for the control group. Error bars indicate SEM. Welch’s t test, two-tailed.

